# Sibling chimerism among microglia in marmosets

**DOI:** 10.1101/2023.10.16.562516

**Authors:** Ricardo C.H. del Rosario, Fenna M. Krienen, Qiangge Zhang, Melissa Goldman, Curtis Mello, Alyssa Lutservitz, Kiku Ichihara, Alec Wysoker, James Nemesh, Guoping Feng, Steven A. McCarroll

**Affiliations:** Department of Genetics, Harvard Medical School, Boston, MA 02115; Stanley Center for Psychiatric Research, Broad Institute of MIT and Harvard, Cambridge, MA 02142, USA; McGovern Institute for Brain Research, Department of Brain and Cognitive Sciences, Massachusetts Institute of Technology, Cambridge, MA 02139, USA; Howard Hughes Medical Institute, Boston, MA 02115

## Abstract

Chimerism happens rarely among most mammals but is common in marmosets and tamarins, a result of fraternal twin or triplet birth patterns in which *in utero* connected circulatory systems (through which stem cells transit) lead to persistent blood chimerism (12-80%) throughout life. The presence of Y-chromosome DNA sequences in organs of female marmosets has long suggested that chimerism might also affect these organs. However, a longstanding question is whether this chimerism is driven by blood-derived cells or involves contributions from other cell types. To address this question, we analyzed single-cell RNA-seq data from blood, liver, kidney and many brain regions across a number of marmosets, using transcribed single nucleotide polymorphisms (SNPs) to identify cells with the sibling’s genome in various cell types within these tissues. Sibling-derived chimerism in all tissues arose entirely from cells of hematopoietic origin (i.e., myeloid and lymphoid lineages). In brain tissue this was reflected as sibling-derived chimerism among microglia (20-52%) and macrophages (18-64%) but not among other resident cell types (neurons, glia or ependymal cells). The percentage of microglia that were sibling-derived showed significant variation across brain regions, even within individual animals, likely reflecting distinct responses by genetic-sibling microglia to local recruitment or proliferation cues or, potentially, distinct clonal expansion histories in different brain areas. In the animals and tissues we analyzed, microglial gene expression profiles bore a much stronger relationship to local/host context than to sibling genetic differences. Naturally occurring marmoset chimerism will provide new ways to recognize the effects of genes, mutations and brain contexts on microglial biology and to distinguish between effects of microglia and other cell types on brain phenotypes.

## Introduction

Chimerism, in which an organism contains cells from genetically distinct animals, happens rarely among mammals. Chimerism is common, however, in the *Callitrichidae* family that consists of the marmosets (*Callithrix*) and their close relatives the tamarins (*Saguinus*): in these primate species, animals usually give birth to dizygotic twins or trizygotic triplets whose blood contains cells from siblings. During development, the siblings share circulation *in utero*, allowing the exchange of hematopoietic stem cells (Gengozian et al., 1969; Wislocki, 1939). Most marmosets then exhibit blood chimerism throughout life: their blood-derived DNA is a mixture of both twins’ genomes, with the twin’s genome contributing 12% to 80% of the DNA in the blood (Niblack et al., 1977; The Marmoset Genome Sequencing and Analysis Consortium, 2014). This indicates that the twins’ hematopoietic stem cells establish permanent residency in one another’s bodies and contribute to blood cell populations throughout life.

A longstanding mystery involves whether other tissues and organs also harbor chimerism. Beyond the blood, Y-chromosome DNA has been detected in the brain and other organs of female marmosets with male twins (Ross et al., 2007; Sweeney et al., 2012), eliciting much speculation about how chimerism might have shaped behavior and natural selection in marmosets. However, it is still not known what cell types harbor this sibling DNA; such observations could in principle be explained by the presence of blood cells within these organs.

Here, we analyze chimerism in the common marmoset (*Callithrix jacchus*) brain, liver, kidney and blood, with single-nucleus RNA-seq (snRNA-seq), using each cell’s RNA-expression pattern to identify its type and using combinations of transcribed SNPs (visible in the snRNA-seq sequence reads) to determine which marmoset sibling is the source of each cell. This approach makes it possible to determine whether chimerism arises from blood cells, resident immune cells, or other cell types, and to explore what chimerism can teach us about cellular migrations and cellular population dynamics.

## Results

### Marmoset chimerism can be characterized at single-cell resolution

To identify which individual cells have the genome of the host marmoset, and which have the genome of the host’s birth sibling, we used combinations of many transcribed SNPs that were visible in the snRNA-seq sequencing reads for each nucleus (using the methods described in Wells et al., 2023).

We first determined whether marmosets have sufficient sequence variation to enable the distinction between host and sibling cells. From whole-genome sequences of 123 marmosets, we identified 13 million polymorphic bi-allelic SNPs in the marmoset genome, with individual marmosets harboring 2.3 to 3.8 million (average 3.4 million) heterozygous sites – comparable to levels of heterozygosity in humans. For sibling comparisons, we detected a large number of sites at which any two siblings’ genomes differed, ranging from 2.0 to 3.7 million sites (average 2.9 million sites) across 96 sibling comparisons (**Fig. 1A**). To determine how many of these sites were visible in snRNA-seq data, we analyzed snRNA-seq data for several marmoset tissues. The results varied by cell type, reflecting that different cell types’ nuclei harbored different quantities of RNA. The hundreds of transcribed variant sites that differed between siblings (median>311 per nucleus) suggested ample power to distinguish between siblings in all cell types (Wells et al., 2023) (**Fig. 1B**).

**Figure 1.**
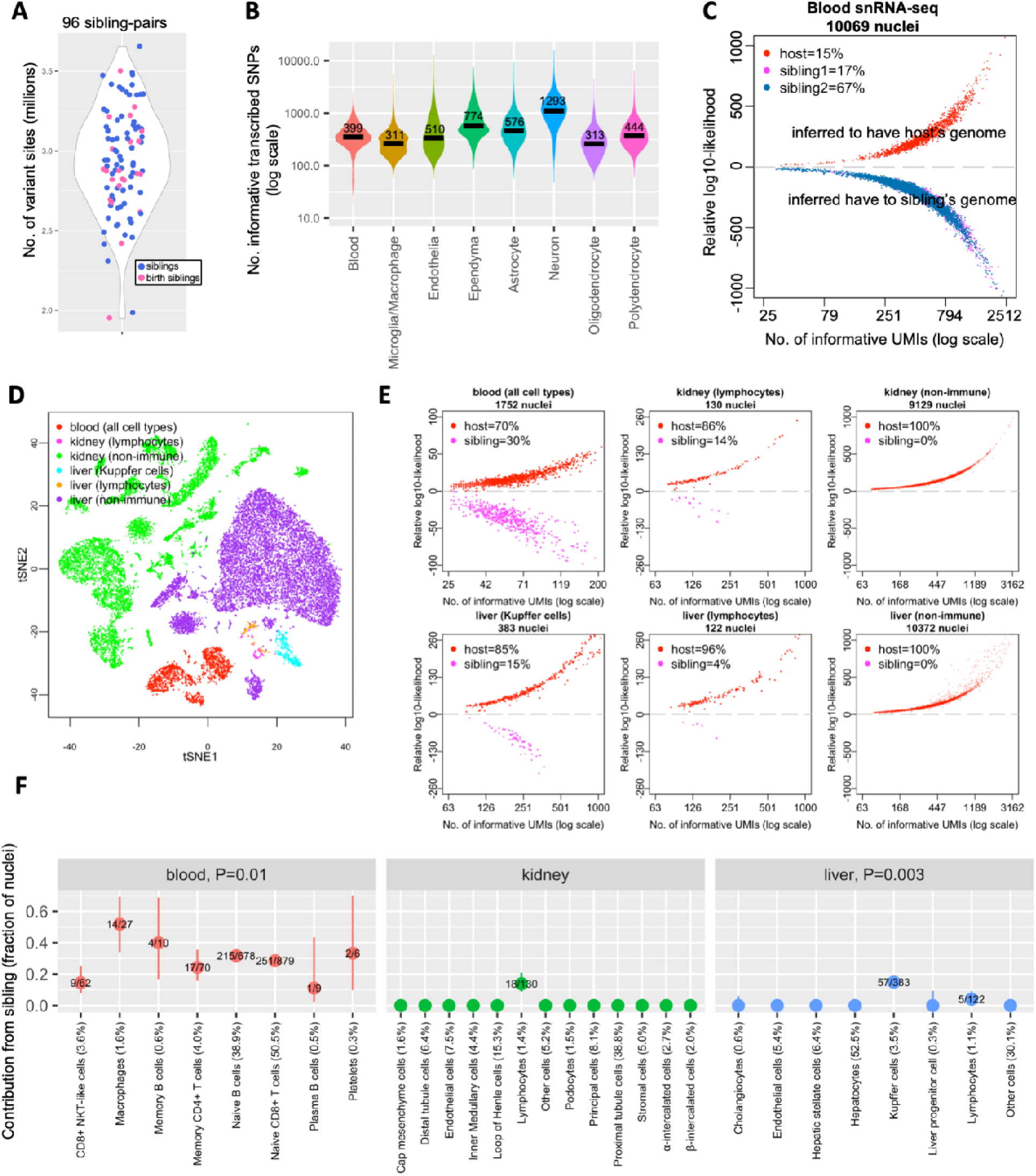
Sibling chimerism analysis at single-cell resolution. **(A)** Numbers of variant sites at which marmoset sibling genomes differ, for 96 sibling-pairs. Each dot is a sibling-pair; birth siblings are colored in pink. **(B)** Numbers of transcribed SNPs (per nucleus) visible in various cell types from blood and brain snRNA-seq datasets of marmoset CJ028. x-axis: snRNA-seq library; y-axis: number of ascertained, transcribed SNPs for which host and sibling have different genotypes; black horizontal lines: median values per cell type. **(C)** Donor-of-origin assignment of each nucleus in blood snRNA-seq of a marmoset. The host marmoset (CJ028) was born with two siblings; each nucleus was assigned to either the host or to one of the two birth siblings. x-axis: number of unique molecular identifiers (UMI; a measure of transcript abundance) that contains SNPs for which the host and sibling’s genomes differ, in log scale; y-axis: inferred likelihood that the cell has host genome minus likelihood that the cell has sibling genome (log10). (**D**) Two-dimensional visualization (tSNE plot) of snRNA-seq data from a marmoset’s (CJ026) blood, kidney and liver. **(E)** Donor-of-origin assignment in marmoset CJ026’s blood, kidney and liver. axes: same as in (C). **(F)** Levels of chimerism in each cell type ascertained in blood, kidney and liver snRNA-seq of marmoset CJ026. y-axis: fraction of sibling cells in each cell type; numbers in fraction: number of sibling nuclei over total nuclei in the cell type; percentage in x-axis labels: cell type representation in the tissue; vertical bars: binomial confidence interval (95%); *P*-values: test of heterogeneity (Chi-square) across immune cell types of a tissue (to test for differences in contribution of sibling across immune cell types).

We next evaluated whether the genome variation visible in snRNA-seq reads was sufficient to distinguish between host and sibling cells. For this we used Dropulation, which identifies the donors of individual cells (from a set of genome-sequenced candidate donors) by using combinations of the transcribed SNPs visible on the snRNA-seq reads of the individual cells (Wells et al., 2023). We first analyzed the blood cells by snRNA-seq of a marmoset (CJ028) born with two birth siblings and used Dropulation (Wells et al., 2023) to assign individual cells to the correct sibling (**Fig. 1C**). The relative likelihoods of the original source of each cell could be strongly differentiated (relative likelihoods of 10^3^ to 10^23^) for >99% of the nuclei (**Fig. 1C**). We found a high level of chimerism: 84% of all nuclei sampled in this marmoset’s blood appeared to contain the genome of one (67% – sibling #1) or the other (17% – sibling #2) of its two birth siblings (**Fig. 1C**), consistent with the wide range of chimerism found in previous studies: 4-82% in marmoset T cells and B cells (Niblack et al., 1977); 13-37% in marmoset whole blood (“The Marmoset Genome Sequencing and Analysis Consortium”, 2014).

### Apparent liver and kidney chimerism arises from infiltrating monocytes

Earlier studies have identified Y-chromosome-derived DNA sequences in the organs of female marmosets with male birth siblings, suggesting that these organs harbor chimerism (Sweeney et al., 2012). However, such observations could also in principle arise from blood or from blood-derived immune cells that are present in those organs (Sweeney et al., 2012).

We performed snRNA-seq analysis of the blood (1,741 nuclei), liver (10,877 nuclei) and kidney (9,262 nuclei) of a marmoset (CJ026) with one birth sibling (**Fig. 1D**). The snRNA-seq profiles clustered into groups that were readily recognized (based on the RNAs expressed) as the principal cell types of each organ; we determined the identity of each cluster using scType, a cell-type identification tool that uses a database of known marker genes (Ianevski et al., 2022).

In kidney and liver, the only clearly twin-derived cells were cells of hematopoietic origin: the resident macrophages in liver (Kupffer cells), lymphocytes in liver, and lymphocytes in kidney (**Fig. 1E,F**). All non-hematopoietic cell types in liver and kidney appeared to contain only the host marmoset’s own genome. Chimerism levels for the two chimeric liver immune cell types appeared to diverge, with sibling-derived cells accounting for 15% of Kupffer cells and just 4% of lymphocytes (5/122 vs 57/383; Chi-square test *P*-value=0.003). In the blood, chimerism levels varied across the various cell types: the most abundant cell types, the Naive B cells and Naive CD8+ T cells, were respectively 29% and 32% sibling-derived, while the less-abundant CD8+ NKT-like cells were 15% sibling-derived (**Fig. 1F**, Chi-square test *P*-value=0.01).

These results indicate that, in this animal’s liver and kidney, apparent DNA chimerism likely arose from infiltrating immune cells rather than other cell types. These results also indicate that cells with siblings’ genomes can differ in their tendency to acquire specific hematopoietic cell fates and in their tendency to infiltrate into organs.

### Marmoset brain microglia and macrophages exhibit abundant chimerism

To characterize chimerism in the marmoset brain, we utilized a large snRNA-seq data set that was generated for a marmoset brain cell atlas (Krienen et al., 2020; Krienen et al., 2023). Brain snRNA-seq was performed on 11 animals (6 adults, 3 neonates and 1 six months old; **Table 1**). All were unrelated except for CJ006 and CJ007 which are birth siblings, and CJ025 and CJ026 which are (non-birth) siblings. All animals come from the three main marmoset colonies that comprise the animals in our facilities: New England Primate Research Center (NEPRC), CLEA Japan, and from a non-clinical contract research organization in Massachusetts. All adult marmosets had no known previous disease and were selected as part of a larger project to create a single-cell atlas of the marmoset brain (Krienen et al., 2020; Krienen et al., 2023). The three neonates died shortly after birth due to unknown reasons and were subsequently selected for snRNA-seq analysis.

**Table 1.**
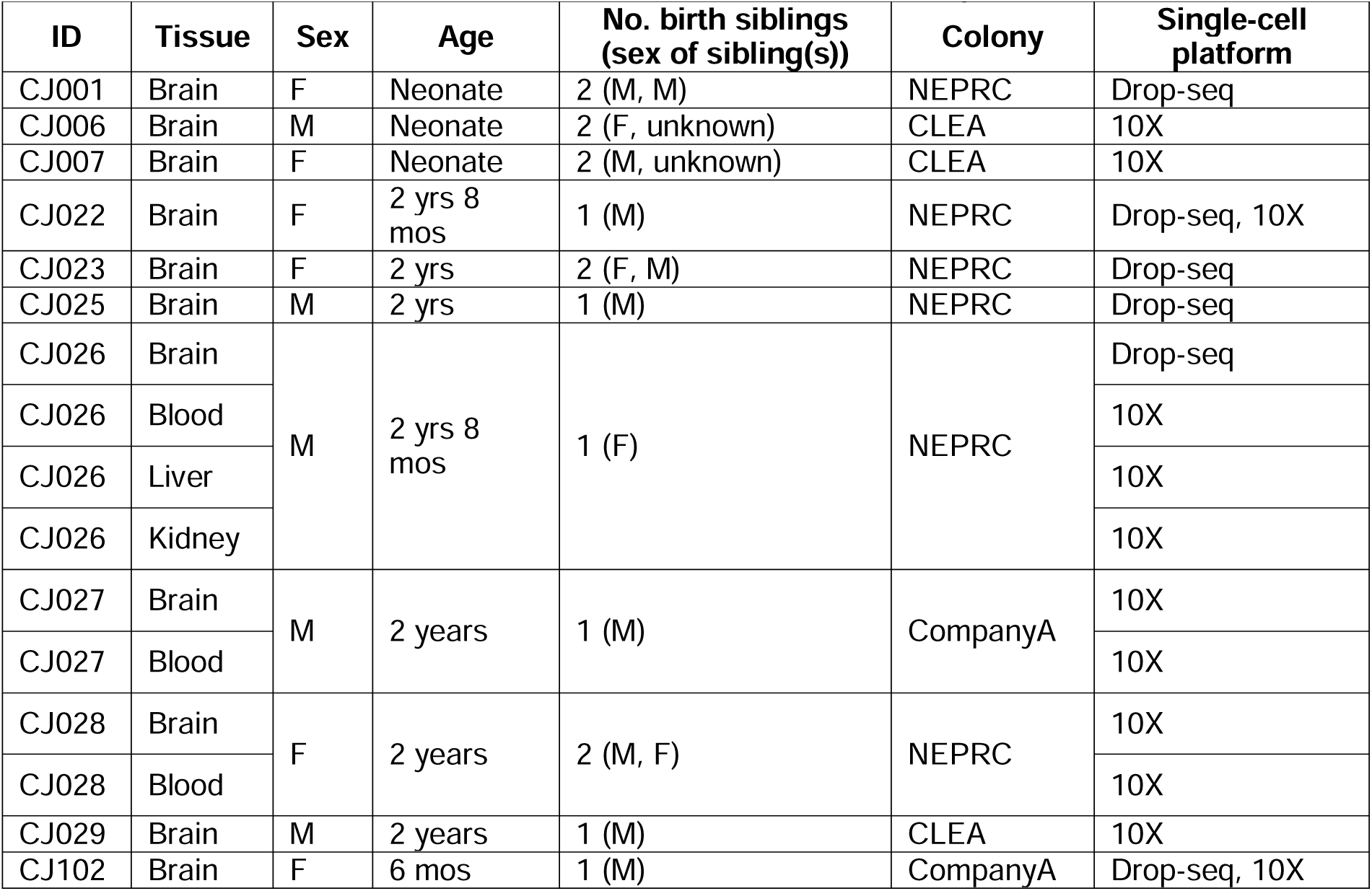
Marmosets analyzed with snRNA-seq in this study. Colonies: NEPRC – New England Primate Research Colony; CLEA – Central Institute for Experimental Animals, Japan; Company A: marmosets obtained from a non-clinical contract research organization.

| ID | Tissue | Sex | Age | No. birth siblings<br>(sex of sibling(s)) | Colony | Single-cell<br>platform |
| --- | --- | --- | --- | --- | --- | --- |
| CJ001 | Brain | F | Neonate | 2 (M, M) | NEPRC | Drop-seq |
| CJ006 | Brain | M | Neonate | 2 (F, unknown) | CLEA | 10X |
| CJ007 | Brain | F | Neonate | 2 (M, unknown) | CLEA | 10X |
| CJ022 | Brain | F | 2 yrs 8<br>mos | 1 (M) | NEPRC | Drop-seq, 10X |
| CJ023 | Brain | F | 2 yrs | 2 (F, M) | NEPRC | Drop-seq |
| CJ025 | Brain | M | 2 yrs | 1 (M) | NEPRC | Drop-seq |
| CJ026 | Brain | M | 2 yrs 8<br>mos | 1 (F) | NEPRC | Drop-seq |
| CJ026 | Blood |  |  |  |  | 10X |
| CJ026 | Liver |  |  |  |  | 10X |
| CJ026 | Kidney |  |  |  |  | 10X |
| CJ027 | Brain | M | 2 years | 1 (M) | CompanyA | 10X |
| CJ027 | Blood |  |  |  |  | 10X |
| CJ028 | Brain | F | 2 years | 2 (M, F) | NEPRC | 10X |
| CJ028 | Blood |  |  |  |  | 10X |
| CJ029 | Brain | M | 2 years | 1 (M) | CLEA | 10X |
| CJ102 | Brain | F | 6 mos | 1 (M) | CompanyA | Drop-seq, 10X |

We first analyzed 497,000 single-nucleus RNA-expression profiles from the neocortex, thalamus, striatum, hippocampus, basal forebrain, hypothalamus and amygdala of an adult marmoset with two birth siblings (marmoset CJ028). We clustered the cell types using gene expression similarities and identified brain cell types as in earlier work (Krienen et al., 2020), identifying neurons, astrocytes, oligodendrocytes, ependymal cells, endothelial cells, microglia and macrophages (**Fig. 2A**). Microglia (which expressed markers *TREM2*, *LAPTM5* and *C3*) and macrophages (which expressed *LYVE1* and *F13A1*) were a small fraction of all nuclei analyzed (about 3.6%), but due to the large number of nuclei we profiled (53 brain tissue dissections, 497,000 nuclei), we were able to ascertain sufficient numbers of microglia (18,185 nuclei) and to a lesser extent macrophages (172 nuclei) for many downstream analyses. We found microglia and macrophages in snRNA-seq data from 10 additional marmosets with different genetic backgrounds from 3 different colonies. Brain snRNA-seq of all 11 marmosets showed consistently the presence of these two myeloid cell types in the brain (**Supplementary Fig. 1**; number of microglia and macrophages in **Supplementary Table 1**).

**Figure 2.**
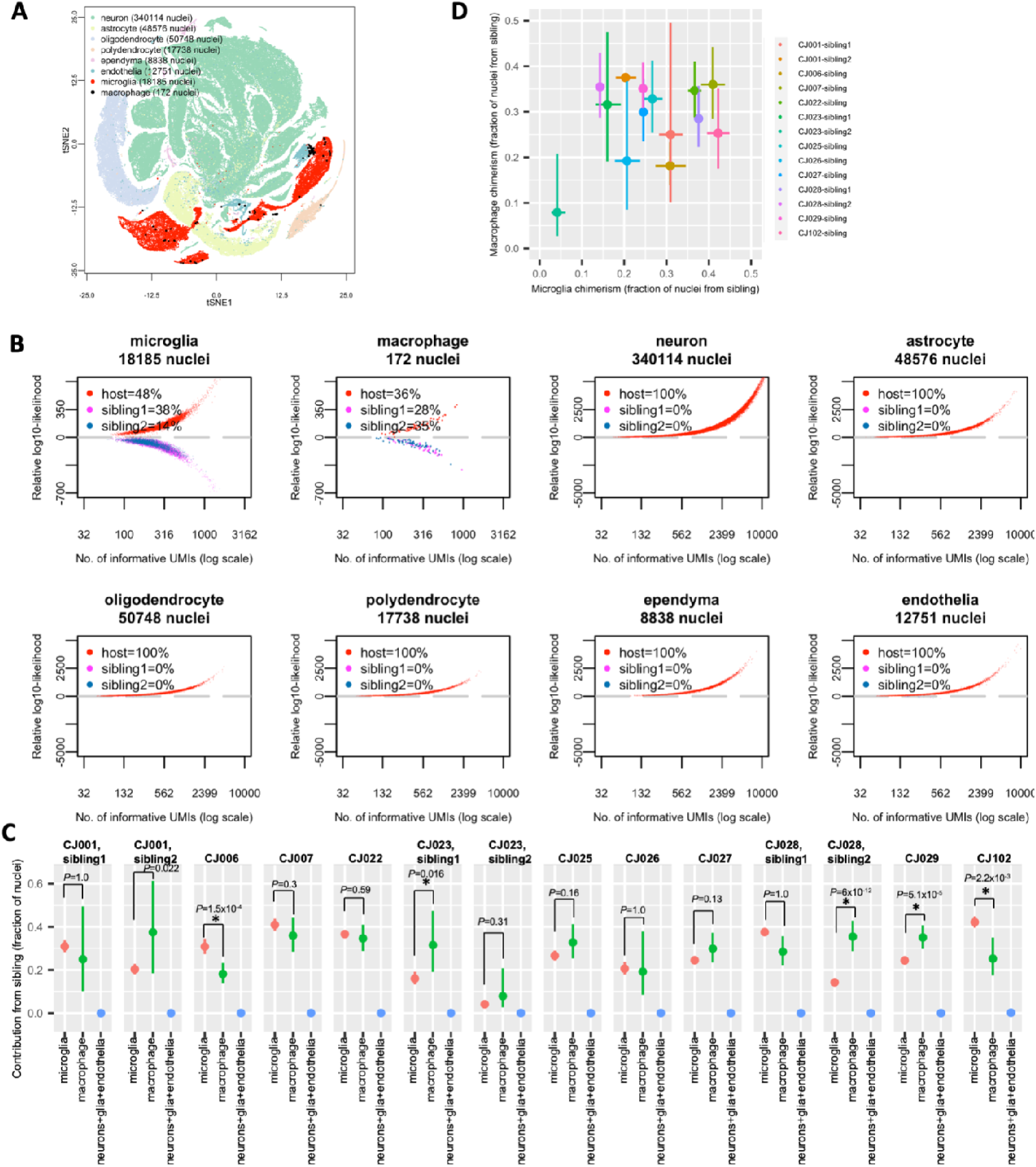
Microglia and macrophages, but not neurons and glia, are chimeric in the marmoset brain. **(A)** Two-dimensional visualization (tSNE plot) of marmoset CJ028’s snRNA-seq profiles from 8 brain regions. **(B)** Donor assignment of each nucleus to one of three possible donors (host, sibling1 or sibling2) of marmoset CJ028’s brain snRNA-seq data. x-axis: number of UMI that contains SNPs for which the host and sibling’s genomes differ, in log10 scale; y-axis: inferred likelihood that the cell has host genome minus likelihood that the cell has sibling genome (log10). Data from all of CJ028’s brain regions. **(C)** Fractions of cells with a sibling’s genome among microglia, macrophages, and other brain cell types (neurons, astrocytes, oligodendrocytes, polydendrocytes, ependyma, endothelial) from 11 marmosets. CJ001, CJ023 and CJ028 were part of a triplet litter and the chimerism fraction of each birth sibling is shown in separate panels. y-axis: fraction of sibling-cells in the cell type. vertical bars: binomial confidence interval (95%). *P*-values: Chi-square test from comparison of chimerism fractions in microglia and macrophage; * (*P*-value<0.05). (**D**) Sibling contributions to microglial (x-axis) and macrophage (y-axis) populations in the same animals. The 14 pairs of animals are from 11 host-sibling1 pairs plus 3 host-sibling2 pairs (three of the 11 host animals were born in a triplet litter; see Table 1). Dots and bars are from (D). Pearson correlation *R*=0.32, 95% confidence interval (−0.26 to 0.73), *P*-value=0.27.

Donor-of-origin analysis of snRNA-seq data from 2.2 million nuclei sampled from 137 brain tissue samples from these 11 marmosets showed a clear and consistent pattern: microglia and macrophages, but not neurons, glia or endothelial cells, harbored chimerism (**Fig. 2B**, **Supplementary Fig. 2**). Microglia exhibited abundant chimerism – across the 11 marmosets, the total fraction of cells with the sibling’s genome ranged from 20% to 52% (for triplets, sum of two siblings; **Fig. 2C**). Macrophages exhibited a similarly wide range of sibling fractions across marmosets (18% to 64%, **Fig. 2C**).

The quantitative extent of microglial chimerism varied across individuals (**Supplementary Fig. 3A**; test of heterogeneity *P*-value<2.2×10^−16^), as did that of macrophage chimerism (**Supplementary Fig. 3B**; test of heterogeneity *P*-value=1×10^−4^). We asked whether microglial and macrophage chimerism were correlated. Intriguingly, only a modest correlation of chimerism levels across 14 host-sibling pairs was observed between the microglia and macrophages (**Fig. 2D**; Pearson correlation 0.31). We investigated further by performing a statistical test that takes into account the uncertainty in the estimates of the chimeric cell proportion using a binomial framework (Methods); in this analysis, microglia chimerism fraction was not a statistically significant predictor of macrophage chimerism fraction (Methods). This suggests that in addition to the cell’s genome, other factors such as local host environment play a role in differential recruitment, proliferation or survival of sibling cells. (We note that macrophages often transit the fluid-filled perivascular space, with a substantially different migration history and arrival dynamics than microglia.) Neither of these two myeloid cell types showed consistently higher chimerism than the other cell type did (**Fig. 2C**).

### Sibling contributions in blood vs. brain

Though microglia (like macrophages) are myeloid cells that derive from hematopoietic stem cells, the ontogenies of microglia and brain macrophages are distinct from those of bone-marrow-derived peripheral blood mononuclear cells (Perdiguero & Geissmann, 2016). As such, differences in the developmental and migration histories of these cell populations could in principle have caused their chimerism fractions to diverge in a systematic way.

We analyzed three marmosets for which snRNA-seq was performed on both blood and brain tissues. Sibling contributions to microglia and brain macrophages were in general quite different from those in blood (**Fig. 3**).

**Figure 3.**
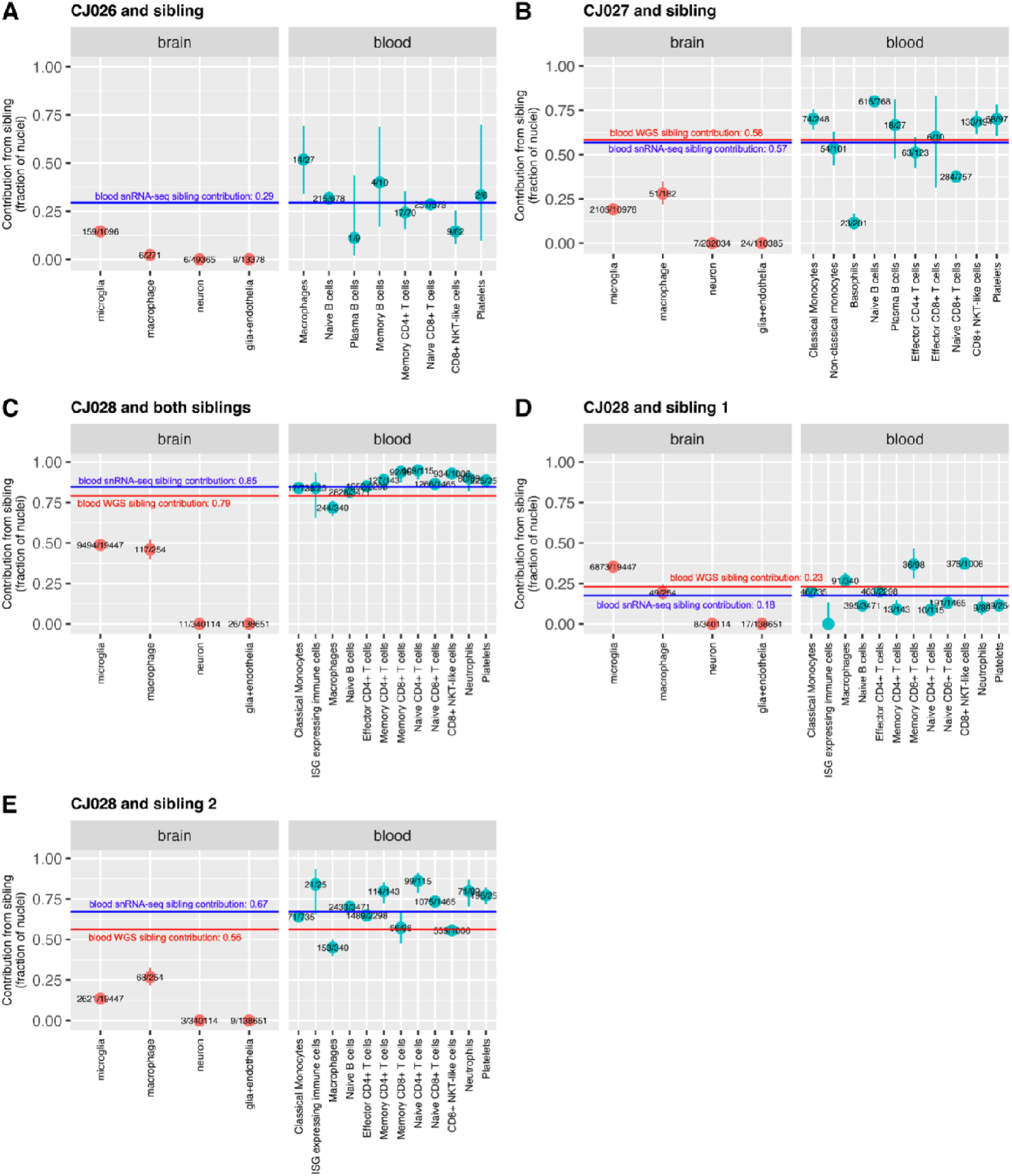
Sibling contributions to hematopoiesis-derived cells diverge between blood and brain. (A)-(E) Chimerism fractions in brain and blood of three animals. CJ028 was part of a triplet litter, and the contribution of each sibling is shown in a separate panel (D,E). Red horizontal lines: twin contribution to blood cells as ascertained from whole-genome sequencing of whole-blood-derived genomic DNA (Census-seq). Blue horizontal lines: total twin contribution to blood cells as estimated from PBMC snRNA-seq (all cells). Vertical bars: binomial confidence interval (95%). Numbers in fraction: number of sibling cells over total cells in the cell type. glia+endothelia: astrocytes, oligodendrocytes, polydendrocytes, ependymal cells, endothelial cells.

Marmoset CJ028’s chimerism (involving two birth siblings) provided a setting in which cells with three different genomes shared the same environment through development until adulthood (to two years of age). Among CJ028’s microglia, the fraction of cells from sibling 1 (35%) was greater than that from sibling 2 (13%) (two-sided test of proportionality *P*-value<2.2^−16^), while in blood, the opposite was true (fraction of cells from sibling 1 across all blood cell types was 18%, fraction of cells from sibling 2 across all blood cell types was 67%; two-sided test of proportionality *P*-value<2.2^−16^) (**Fig. 3D,E**, **Supplementary Table 2**).

### Microglia chimerism fraction varies across brain regions

Sibling contributions to the microglial population could in principle be shaped by effects that are local to specific brain areas, including differential response of sibling microglia to local recruitment or proliferation cues, or population bottlenecks such as clonal expansions. To evaluate whether the sibling contribution to the microglial population varied across brain areas within individual marmosets, we performed chimerism analysis for each of the brain regions profiled in the snRNA-seq datasets: neocortex, thalamus, striatum, hippocampus, basal forebrain, hypothalamus and amygdala (Krienen et al., 2020; Krienen et al., 2023). Within each marmoset, the fraction of microglia with a sibling’s genome diverged across a marmoset’s brain regions (**Fig. 4**). For example, in marmoset CJ025, sibling contributions to microglial populations ranged from 11% (21/193) in the thalamus to 56% (174/310) in the striatum (*P*-value=1.1×10^−23,^ Chi-square test of thalamus vs striatum; *P*-value=1.5×10^−40^, Chi-square test across all 4 brain regions). For marmoset brains profiled with at least 300,000 nuclei, CJ027, CJ028 and CJ029, tests of heterogeneity *P*-values were even more significant: 8.8×10^−83^, <1×10^−300^, and 6.1×10^−47^, respectively (the error bars are very short in the corresponding panels in Fig. 4). We used the binomial generalized linear mixed-model framework and found that all brain regions were statistically significant predictors for microglia chimerism fraction, supporting the conclusion that chimerism varies across brain regions (Methods). Analysis of finer brain substructures showed a similar result **(Supplementary Fig. 4**; the binomial generalized linear mixed-model framework determined that 18 out of 27 brain substructures were statistically significant as predictors for microglia chimerism fraction, Methods). None of the brain regions exhibited consistently higher or lower chimerism levels, suggesting that these divergences did not result from differential physical access of host and sibling microglia to different brain areas (**Fig. 4** and **Supplementary Fig. 4**).

**Figure 4.**
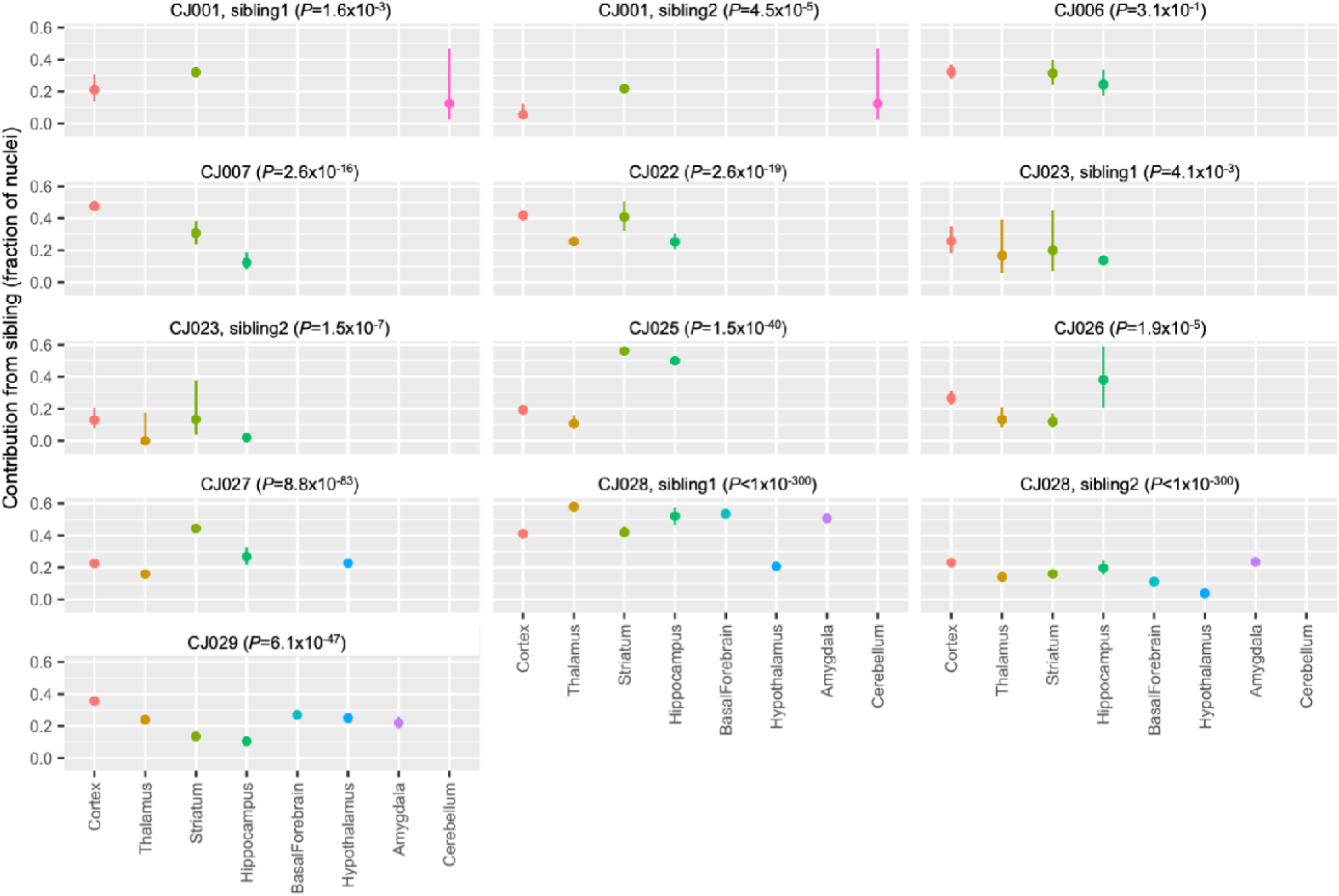
Sibling contributions to brain microglial populations vary across an animal’s brain areas. Contributions of sibling(s) to the microglial populations ascertained in principal brain areas. CJ001, CJ023 and CJ028 were part of a triplet litter and the chimerism contribution of each twin is shown in a separate panel. CJ102 was profiled in only one brain region and hence was not included in the analysis. Brain regions with missing data were not profiled in that animal. y-axis: fraction of twin cells. Vertical bars: binomial confidence interval (95%); *P*-values: test of heterogeneity across an animal’s brain regions.

### Gene expression comparisons of host– to sibling-derived microglia

Chimerism provides the unusual opportunity to compare cells with different genomes in a shared *in vivo* biological context. We compared RNA expression between host– and sibling-derived microglia of a female marmoset with two birth siblings. Sex differences among the siblings (the host (CJ028) was a female and one of the two siblings was a male) allowed a natural control: the *XIST* gene encodes a non-coding RNA involved in silencing one copy of chromosome X in females and thus exhibits sex-specific expression due to cell-autonomous mechanisms. We found that (as expected) *XIST* transcripts were detected at far higher levels in the snRNA-seq profiles of microglia with the female twin’s genome relative to the male twin’s (**Fig. 5A,C**). By contrast, *XIST* transcripts were detected at similar levels in two microglial populations with the genomes of female twins (**Fig. 5B**).

**Figure 5.**
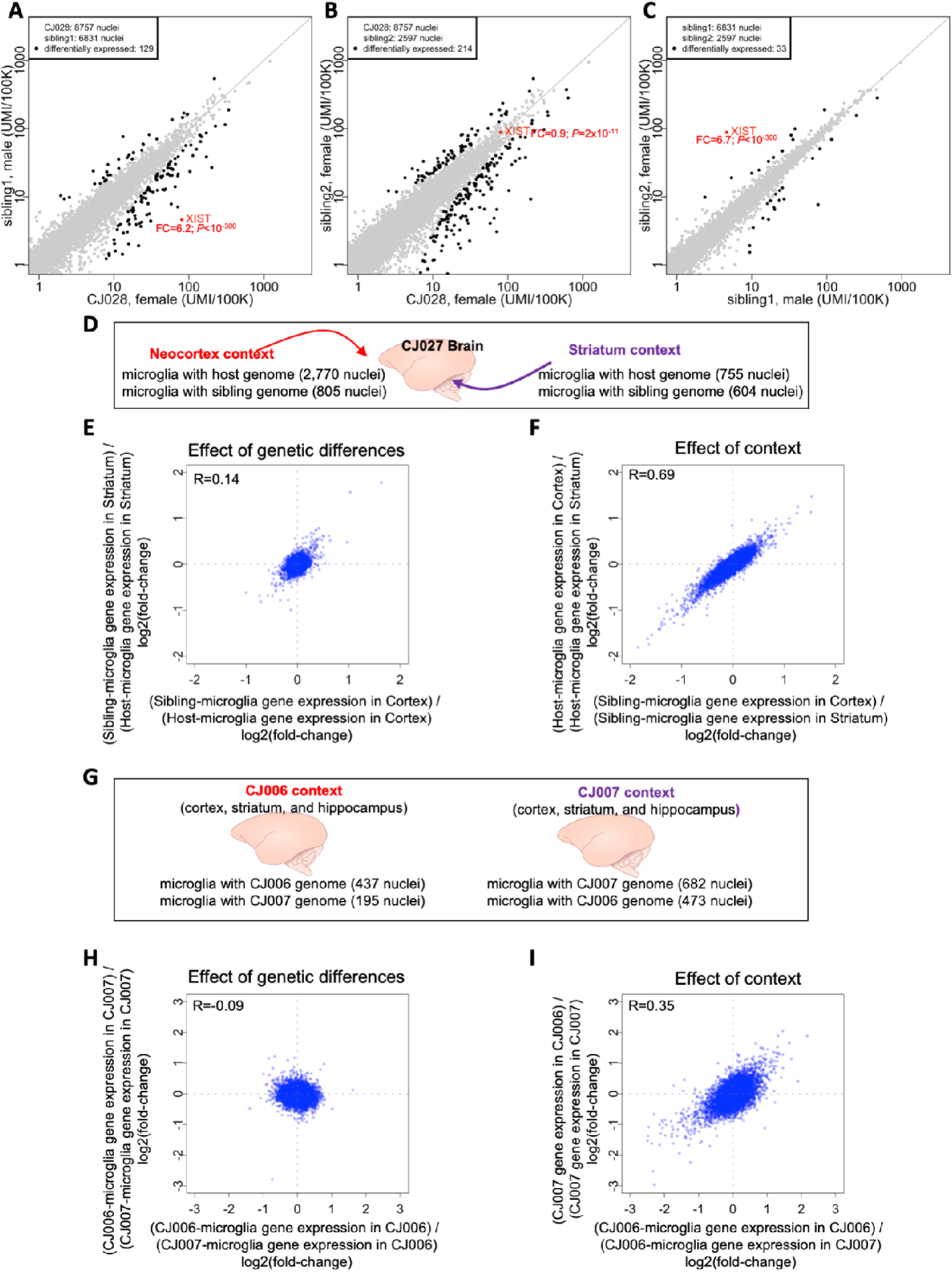
Utilizing natural chimerism to distinguish cell-autonomous from non-cell-autonomous effects on gene expression, and to compare the effects context and genetic variation in shaping gene expression. (A-C) Comparisons of RNA expression between microglial populations within host animal CJ028, who had two birth siblings. Comparisons of gene expression between microglia with the genomes of **(A)** the female host and male sibling, **(B)** the female host and female sibling, and **(C)** the two siblings (male and female). In (A) to (C), each point represents a gene; its location on the plot represents the level of expression of that gene among microglia with two different genomes in the same animal. x– and y-axes: normalized gene expression levels (number of transcripts per 100,000 transcripts). FC: fold-change of gene expression, female/male for *XIST*. Fold-change and *P*-values were calculated using the binomTest method from the edgeR package (Robinson et al., 2010). Differentially expressed genes (black dots) were defined as: FDR *Q*-value<0.05 and fold-change>1.5 (in either direction) and the gene must be expressed in at least 10% of at least one of the two sets of microglia being compared. **(D-I)** Higher effect of context than genetic differences in shaping gene expression. **(D)** In the brain of marmoset CJ027, the neocortex and striatum are two contexts where two sets of microglia with different genomes reside. **(E)** Effect of genetic differences. x-axis: log2-fold-change of cortical microglia gene expression between host and sibling cells; y-axis: log2-fold-change of striatal microglia gene expression between host and sibling cells. **(F)** Effect of context. x-axis: log2-fold-change of the sibling’s gene expression between the two brain regions; y-axis: log2-fold-change of the host’s gene expression between the two brain regions. **(G-I)** The brains (cortex, striatum and hippocampus) of two birth siblings provide biological contexts in which populations of microglia with two sibling genomes reside. The effect of genetic differences (H) and effect of animal context (I) are compared, for the same brain areas (combined data from cortex, striatum and hippocampus). x– and y-axes: log2-fold-changes of the gene expression between two sets of microglia (the sets being compared are indicated in the axis labels). *R*: Spearman correlation.

Gene-expression differences between host– and sibling-derived microglia in the same brain could in principle arise from asymmetries in their developmental histories (which would be shared across host animals) or from genomic differences (which would vary from host animal to host animal). In all eleven individual marmosets, analysis identified genes whose differential expression distinguished microglia with the two sibling genomes (hundreds of genes in total), documenting a substantial effect of sibling genetic differences on microglial gene expression. However, we did not find any gene whose expression level recurrently distinguished “host” microglia (microglia with the same genome as neural cell types) from “guest” microglia (microglia with the sibling genome), aside from the XIST gene (a proxy for sibling sex differences, which were of course common) (**Supplementary Fig. 5**, **Fig. 5A-C**). In other words, although there were always gene-expression differences between sibling microglia, none of them consistently distinguished between host and guest microglia, suggesting that they were instead due to sibling genetic differences. We note that both analyses are power-limited, as the number of microglia in most animals, especially guest microglia, were modest (**Supplementary Fig. 5**); thus, we cannot rule out the possibility that there may be one or more genes whose expression levels reflect developmental histories (host vs. guest origin), just as there are likely far more genes (than the hundreds we identified) that can have sibling expression differences due e.g. to genetic differences between siblings. We sought to increase power (beyond single-gene analysis) by using latent factor analysis (Ling et al., 2024) to identify and quantify the expression of microglial gene-expression programs; however, even this analysis did not find any gene expression programs that exhibited consistent host-twin differences in expression levels (Methods).

### Brain context vs. genetic differences as determinants of microglial gene expression

Chimerism presents interesting opportunities to distinguish between cell-autonomous and contextual effects on a cell’s biology, and to compare the magnitudes of such effects.

We first considered the difference in contexts provided by pairs of brain areas by analyzing snRNA-seq data from the neocortex and striatum of marmoset CJ027; the resident microglial populations with different genomes make it possible to compare contextual to genetic effects on microglial gene expression (**Fig. 5D**). Genetic effects appeared to elicit very many small-magnitude gene-expression differences; these differences were shared between cortical and striatal microglia (**Fig. 5E**). Brain-area context elicited much larger-magnitude gene-expression differences, which were experienced in common by microglia with both genotypes (**Fig. 5F**). We obtained similar results for all pairs of brain areas analyzed (52 context vs genetic effect from brain snRNA-seq of 6 marmosets with at least 60 cells available for analysis in each context; **Supplementary Table 3**; **Supplementary Fig. 6**).

We next considered the difference in contexts provided by the same brain area in different marmosets. Two of the marmosets profiled, CJ006 and CJ007, were birth siblings who passed away as neonates (the only birth siblings in our dataset), and thus provided the additional opportunity to distinguish genetic from contextual effects by analyzing the two sibling microglial populations in the cortex, striatum and hippocampus of both marmosets (**Fig. 5G**). The effects of context (host marmoset) in microglia from all three brain areas appeared to be far larger than the cell-autonomous effects of genetic differences (**Fig. 5H,I**).

## Discussion

A longstanding debate concerns the extent of chimerism in marmosets and tamarins. Chimerism in these species has been detected in diverse organs but arises from unknown cell types (Ross et al., 2007; Sweeney et al., 2012). Here we found that chimerism in the brain, liver and kidney is present but appears to arise entirely from cells of the myeloid and lymphoid lineages, including infiltrating macrophages, monocytes, and microglia.

Cells of the myeloid and lymphoid lineages derive developmentally from hematopoietic stem cells. We found no strong evidence of chimerism among 2.2 million non-hematopoietic cells in the liver (from one marmoset), kidney (from one marmoset) or brain (from 11 marmosets). Thus, while marmosets share a circulation *in utero*, we found no evidence that other kinds of stem cells or progenitors (beyond those of hematopoietic lineage) had been shared via this route in any appreciable number. However, we found that in the marmoset brain, the microglia and macrophages, which also derive from this lineage, routinely harbor abundant chimerism, with 10-50% of a marmoset’s microglia containing the genome(s) of birth sibling(s).

Organs in the same marmoset (liver, kidney, brain) differed markedly in the sibling contribution to resident macrophage and monocyte populations, with microglial chimerism fraction (the fraction of cells contributed by siblings) varying by as much as 40 percentage points across a marmoset’s brain areas. This phenomenon has more than one potential explanation. First, cells from the host and sibling could in principle respond differently to recruitment or proliferation cues that vary spatially; if this is the case, marmoset chimerism could provide a model for studying the effects of mutations and natural sequence variation on cell migration and recruitment. (Although we found that genetic effects were smaller than contextual effects in shaping microglial gene expression at any moment in time, genetic effects were clear (**Fig. 5E**), and even small effects on proliferation rates would tend to have effects that increase exponentially over time.) Second, beyond such recruitment effects, it is also possible that these differences suggest a substantial role of clonal expansions and population bottlenecks in shaping local microglial and macrophage populations.

We found that the cellular contribution of birth siblings to myeloid cell populations was significantly different in blood than in brain in the modest number of marmosets analyzed (**Fig. 3**). Unlike the differences among brain areas, the blood-brain differences tended to be directional, with more-modest sibling contributions in the brain than in the blood. Though this would need to be confirmed in many more marmosets to be definitive on its own, it is plausibly connected to this aspect of marmoset fetal development: sharing of a blood circulation between the two fetuses occurs during a window that is more temporally extended than the waves of colonization of the brain by microglia, potentially allowing for greater exchange in the centers of blood hematopoiesis (the liver and then the bone marrow). In microglia and macrophages, due to the defined waves of hematopoiesis in the yolk sac and migration patterns of microglia and macrophages to the developing brain, the opportunity for a progenitor cell from a twin to colonize a host’s brain may need to occur during a more restricted temporal window.

Comparisons of gene expression between microglial cells with host and sibling genomes in a shared brain context may provide many future opportunities to distinguish the cell-autonomous from non-cell autonomous genetic effects of genetic differences and engineered mutations. Such analyses could become especially useful scientifically as genome editing increasingly enables the utilization of marmosets as a model organism in translational neuroscience (Aida & Feng, 2020; Feng et al., 2020). Our pilot analysis of host-sibling microglial gene expression differences in the brains of two co-twins revealed a large role of animal context (relative to genetic differences) in shaping microglial gene expression. This result points to an important principle: the ability to isolate the effects of a mutation will be greatly strengthened by the ability to make within-animal (rather than just between-animal) comparisons of cells with different genotypes.

A long history of innovation in genetics involves elaborate ways to create mosaics in mice, *C. elegans* and other laboratory organisms in order to distinguish cell-autonomous from non-cell autonomous genetic effects. Natural chimerism in marmosets may enable many straightforward ways to pursue such kinds of studies. Natural chimerism may also make it possible to determine when microglia or macrophages, as opposed to other cell types, mediate the effect of a mutation on an animal’s phenotype.

Chimerism could also enable interesting future analyses of whether there are adaptive benefits of chimerism in marmoset immune cells, among whom chimerism could in principle allow presentation of a wider variety of antigens for adaptive immunity. In a recent outbreak of yellow fever in Brazil in 2016-2018, marmosets were found to be less susceptible than other primates that lack immune system chimerism, including the howler monkeys (*Alouatta*), robust capuchins (*Sapajus*), and titi monkeys (*Callicebus*) (de Azebedo Fernandes, et al., 2021). In studying future outbreaks in marmosets, one could use single-cell RNA-seq and the methods described here to study how genetically distinct immune cells (in the same animal) have differentially migrated to affected tissues and/or assumed “activated” immune cell states. Recent innovations in spatial transcriptomics with sequencing readouts (that detect SNP alleles) may also make it possible to identify any differential recruitment of genetically distinct immune cells to focal infection sites.

Microglia perform essential roles in the development and regulation of the central nervous system (CNS), including by sculpting or “pruning” neuronal circuits (Hammond et al., 2018; Schafer et al., 2012; Schafer & Stevens, 2015), and are implicated in or hypothesized to contribute to a wide range of brain disorders and diseases, including Alzheimer’s disease, Parkinson’s disease, autism spectrum disorder, and schizophrenia. Marmoset microglial chimerism will enable many new ways of studying microglia and the effects of genes and alleles upon brain biology.

## Methods

### Ethical compliance

Marmoset experiments were approved by and in accordance with Massachusetts Institute of Technology IACUC protocol number 051705020.

### Nucleus Drop-seq library preparation and sequencing

Nucleus suspensions were prepared from frozen tissue and used for nucleus Drop-seq following the protocol we have described at https://doi.org/10.17504/protocols.io.2srged6. Drop-seq libraries were prepared as previously described (Macosko et al., 2015), with modifications, quantification and quality control as described in a previous study (Saunders et al., 2018), as well as the following modifications optimized for nuclei: in the Drop-seq lysis buffer, 8M guanidine hydrochloride (pH 8.5) was substituted for water, nuclei were loaded into the syringe at a concentration of 176 nuclei/μl, and cDNA amplification was performed using around 6,000 beads per reaction, 15 PCR cycles. Raw sequencing reads were aligned to the calJac3 marmoset reference genome assembly and reads that mapped to exons or introns of each assembly were assigned to annotated genes (https://github.com/broadinstitute/Drop-seq). Drop-seq libraries are indicated in **Table 1**.

### Nucleus 10X Chromium library preparation and sequencing

Single-nucleus suspensions from frozen tissue were generated as for Drop-seq; GEM generation and library preparation followed the manufacturer’s protocol (protocol versions #CG00052 Chromium Single Cell 3’ v2 and #CG000183 Chromium Single Cell3′ v3 UG_Rev-A). Raw sequencing reads were processed and aligned using the same method for aligning Drop-seq reads. 10X Chromium libraries are indicated in **Table 1**.

### Clustering of cells using Independent Component Analysis

Nuclei from intact cells were identified and clustered into cell types using a method that we have previously described (Krienen et al., 2020; Saunders et al., 2018). Briefly, nuclei with less than 400 detected genes were not used in the analysis. A digital gene expression matrix was created for a set of libraries from the same animal that were to be co-analyzed (**Supplementary Table 4**), and independent component analysis using the fastICA package in R was used after normalization and variable gene selection as previously described (Krienen et al., 2020; Saunders et al., 2018). A Louvain-based clustering algorithm was performed on the top 60 independent components. Due to the large number of nuclei profiled in some marmosets (CJ027, CJ028, CJ029), memory requirements exceeded machine limits and for these marmosets, we divided the clustering analysis into two or three batches (**Supplementary Table4**). The brain of marmoset CJ022 was profiled using both Drop-seq and 10X and a separate clustering was done for each snRNA-seq method (**Supplementary Table 4**). We ran the clustering algorithm 12 times using 3 nearest neighbor parameters (10,20,30) and 4 resolution parameters (0.3, 0.5, 0.1, 1.0). Markers for each cluster were identified using differential gene expression analysis (Krienen et al., 2020; Saunders et al., 2018). We inspected each clustering result and chose the one which yielded separate clusters for microglia and macrophages (**Supplementary Table 4**).

### Identification of cell types

For the brain datasets, the microglia and macrophage clusters were identified by the markers *TREM2, C3, LAPTM5* for microglia and *F13A1*, *LYVE1* for macrophages. The other brain cell types were identified using cell type markers for neurons, astrocytes, oligodendrocytes, polydendrocytes and endothelial cells that we used as before (Krienen et al., 2020). Cell types in blood, liver and kidney were identified using the ScType method (Ianevski et al., 2022).

### Donor-of-origin analysis and detection of host-sibling doublets (Dropulation)

We used the Dropulation suite to calculate a donor likelihood for each cell (Wells et al., 2023) (software available at https://github.com/broadinstitute/Drop-seq). The host and birth sibling genotypes were provided as input to Dropulation’s AssignCellsToSamples tool, together with the snRNA-seq BAM file and a list of cell barcodes that were identified to be intact cells. To generate chimerism-free reference genotypes, we cultured fibroblasts and performed whole genome sequencing (WGS) on the resulting DNA. We found that the difference in likelihoods between host and sibling increase with the number of UMI of the cell, and hence we imposed a minimum number of UMI for each marmoset’s cells (**Supplementary Table 4**). We also performed doublet detection using Dropulation’s DetectDoublets to obtain a likelihood of a cell having a mix of transcripts from the host and sibling(s). Doublets lie between the host and sibling curves (**Supplementary Fig. 7**) and for each marmoset we empirically obtained a threshold for the Dropulation test statistic to identify them and were discarded in all analyses (**Supplementary Fig. 7**, **Supplementary Table 4**). The likelihoods plotted in Figures 1C,1E,2C and Supplementary Fig. 2 are from cells that have been filtered for minimum UMI and doublets.

Marmosets CJ006 and CJ007 were born in a triplet litter (tri-zygotic) that all died shortly after birth. We did not have access to any tissue from the third sibling and were not able to perform whole genome sequencing on it. For Dropulation analysis of CJ006 and CJ007’s brains, we provided only the genotypes of marmosets CJ006 and CJ007. Nuclei that contain the genome of the third unknown sibling will mostly be identified as doublets, which were discarded in our analysis.

### Additional filtering for microglia and macrophage clusters

We performed additional filtering of microglia and macrophage cells. When we compared the gene expression of host microglia and sibling microglia using cell-types from first-round clustering (and with UMI and doublet filtering), we found an abundance of genes that have higher expression in host than in the sibling (**Supplementary Fig. 8**, see panels B, C, G, I and K). The asymmetry could arise from neuronal cells mis-classified as microglia or macrophages. To filter out these mis-classified cells, we sub-clustered the microglia and macrophage cell types of each marmoset using the same fastICA and Louvain-based clustering used in the first round of clustering. We found that some sub-clusters were not chimeric, indicating that they were not cells of hematopoietic origin, and that discarding these cells improved the symmetry between host and sibling gene expression (**Supplementary Fig. 8**).

### Whole genome sequencing

Illumina libraries from fibroblast, blood, brain and buccal cells (**Supplementary Table 5**) were created as follows. An aliquot of genomic DNA (150ng in 50μL) is used as the input into DNA fragmentation (aka shearing). Shearing is performed acoustically using a Covaris focused-ultrasonicator, targeting 385bp fragments. Following fragmentation, additional size selection is performed using a SPRI cleanup. Library preparation is performed using a commercially available kit provided by KAPA Biosystems (KAPA Hyper Prep with Library Amplification Primer Mix, product KK8504), and with palindromic forked adapters using unique 8-base index sequences embedded within the adapter (purchased from Roche). The libraries are then amplified by 10 cycles of PCR. Following sample preparation, libraries are quantified using quantitative PCR (kit purchased from KAPA Biosystems) with probes specific to the ends of the adapters. This assay is automated using Agilent’s Bravo liquid handling platform. Based on qPCR quantification, libraries are normalized to 2.2nM and pooled into 24-plexes. Sample pools are combined with NovaSeq Cluster Amp Reagents DPX1, DPX2 and DPX3 and loaded into single lanes of a NovaSeq 6000 S4 flowcell cell using the Hamilton Starlet Liquid Handling system. Cluster amplification and sequencing occur on NovaSeq 6000 Instruments utilizing sequencing-by-synthesis kits to produce 151bp paired-end reads. Output from Illumina software is processed by the Picard data-processing pipeline to yield CRAM or BAM files containing demultiplexed, aggregated aligned reads. All sample information tracking is performed by automated LIMS messaging. All samples were sequenced to 30X coverage.

### Variant site detection and genotyping from whole genome sequencing

Illumina paired-end reads were aligned to the calJac3 reference marmoset genome assembly using bwa (Li & Durbin, 2010) with command “bwa mem”. Duplicate reads were marked using Picard Markduplicates and for each chromosome, the GATK Haplotype Caller (McKenna et al., 2010) was run in genotype discovery GVCF mode. For each chromosome, the GVCFs of all samples analyzed in this study (from fibroblasts, blood, buccal cells, skin, brain and hair), were combined into a single GVCF file using GATK CombineGVCFs. To obtain the highest sensitivity in calling SNPs, we included in the GVCF additional fibroblasts whole genome sequences from the colony, yielding a total of 113 marmosets for multi-sample variant calling. The GVCF of each chromosome was genotyped using GATK GenotypeGVCFs. Only bi-allelic SNPs were used in the analysis and the following filters were used: QD<4.0 | FS>60.0 | MQ<40.0 | MQRankSum<-12.5 | ReadPosRankSum<-8.0 | MAF<0.01 | QUAL<500. SNP calls from all chromosomes were combined into one VCF file and additional filtering was performed to discard heterozygous sites that exhibited extreme allelic imbalance, i.e., the fraction of non-reference allele (from all samples) is less than 0.2 or greater than 0.8, and furthermore, sites in copy number variant regions were discarded (copy number variant regions were obtained by running Genome STRiP (Handsaker et al., 2015) on whole genome sequencing data from 113 fibroblast samples).

### Dropulation analysis using sibling genotypes from whole genome sequencing of buccal cells

For four marmosets in our dataset (CJ022, CJ025, CJ026, CJ102; all born with one sibling and the siblings are CJ106, CJ104, CJ105 and CJ103, respectively), only the buccal cells (from cheek swabs) of their siblings were available for whole genome sequencing. Using a method that quantifies chimerism from whole genome sequencing data (Census-seq; software available at https://github.com/broadinstitute/Drop-seq) (Mitchell et al., 2020), we estimated the chimerism fraction in buccal cells as follows: CJ106: 10%, CJ104: 24%, CJ105: 24% and CJ103: 9%. Thus, the genotypes we obtained for these marmosets will include errors and those genotyping errors could subsequently affect the Dropulation (donor-of-origin) analysis that was used to estimate chimerism. To empirically estimate how sibling genotypes obtained from a chimeric tissue affect Dropulation analysis, we selected a host-sibling pair whose genome sequencing were both obtained from fibroblast cultures: CJ027 and its birth sibling CJ140. To simulate DNA contamination, we fixed the sequencing coverage of CJ140 to 40X, and replaced between 1% to 60% of the reads from CJ027’s sequencing reads (random subsampling using ‘samtools view –s’). We genotyped CJ140’s “chimeric” bam files using GATK’s “genotype given alleles” mode and compared the genotypes with CJ140’s true genotypes from its pure fibroblast WGS. We found that the sensitivity at heterozygous sites remains constant with different contamination levels, while the false positive rate increases. The false positive calls at heterozygous sites come from homozygous sites incorrectly genotyped as heterozygous (**Supplementary Fig. 9A-F**). Next, we re-analyzed donor-of-origin on brain snRNA-seq of CJ027 sibling (CJ140) genotypes from simulated contaminated DNA (300,000 nuclei; CJ027 genotypes from pure fibroblast WGS, CJ140 genotypes from WGS with various contamination levels). We found that chimerism in CJ140’s WGS resulted in doublets being assigned to the twin (**Supplementary Fig. 9G-L**), which subsequently causes a slight increase in chimerism estimates (**Supplementary Fig. 9M-U**). The contamination levels in buccal cells were from 9% to 24%, which we estimate will result in an overestimation in microglia chimerism of up to 3.5 percentage points and 4.5 percentage points in macrophage chimerism. Our results will not be affected by this over-estimation of chimerism in 4 marmosets since the conclusions were made from the analysis of all 11 marmosets (including 7 marmosets whose siblings were genotyped from fibroblast cultures).

### Gene expression analysis of host and sibling meta cells

For each cell type, the host and sibling “meta cells” were calculated from the sum of UMI counts per gene across cells and were scaled to counts per 100,000 transcripts. The fold-changes and *P*-values of differentially expressed genes were identified using the binomTest method from the edgeR package (Robinson et al., 2010). The filters we used to identify statistically significant differentially expressed genes are: FDR Q-value<0.05 and fold-change>1.5 (in either direction) and the gene must be expressed in at least 10% of at least one of the two sets of microglia being compared.

### Binomial generalized linear mixed-effects model analysis

To perform an analysis of **Fig. 2D** that takes into account the uncertainty in the estimate of the chimeric cell proportion, we performed a binomial generalized linear mixed-effects model analysis in R using the command **glmer(y∼(1|indiv)+ chimerism_micro, family=binomial)**, where **y** is a vector (of length 1,333) containing the genomic identity of each macrophage (either host or twin), **1|indiv** models a random effect for the identity of each animal, and **chimerism_micro** is the microglia chimerism of the animal’s brain. The fixed effects probability of **chimerism_micro** was 0.795 indicating that microglial chimerism fraction was not statistically significant as a predictor for macrophage chimerism fraction. The estimate for the intercept was –0.8115 and estimate for **chimerism_micro** was 0.3106, which indicates that the probability of a cell is a macrophage given the microglia chimerism fraction was only 0.57 (plogis(−0.8115+0.3106)).

We used the same framework to further analyze **Fig. 4**. We included brain region as a covariate in the binomial framework: **glmer(y∼(1|indiv) + brain_reg + assay, family=binomial)**, where **y** is a vector of length 48,439 containing the genomic identity of each microglia (either host or twin) and **assay** is either “Drop-seq” or “10X”. The brain regions assayed in **Fig. 4** are the cortex, hippocampus, hypothalamus, striatum, thalamus, and basal forebrain. All these brain regions were statistically significant as predictors for microglia chimerism fractions (all *P*-values<2×10^−16^), supporting the conclusion that chimerism varies across brain regions. We also re-analyzed **Supplementary Fig. 4** using the same framework and found that 18 out of 27 brain substructures were statistically significant as predictors for microglia chimerism fraction.

### Latent factor analysis

Following the method described in (Ling et al., 2024), we performed latent factor analysis using the probabilistic estimation of expression residuals (PEER, Stegle et al., 2010) on the gene-by-donor matrix expression of microglia. We started by creating a gene-by-cell matrix of microglia gene expression from all animals and we normalized the matrix using SCT transform version 2 (Choudhary and Satija, 2022) with 3000 variable features. We obtained the Pearson residuals from SCT normalization and summed up the residual across cells with the same genome to obtain a gene-by-donor matrix of expression measurements of microglia. We used this matrix as input to PEER and ran the tool with a provided number of factors from 9 to 12. For each gene expression latent factor, to evaluate whether host/sibling identity has a consistent effect on expression levels, we performed a linear regression with host/sibling identity using **glm(peer_factor_k ∼ host_or_twin)**. For all factors, the *P*-values for the effect of **host_or_twin** were all insignificant (greater than 0.1), indicating that no PEER factor associated with host-vs-twin identify. Thus our results found no large-scale gene expression program that was consistently expressed differently between hosts and twins.

### Software availability

All software used in the analysis are publicly available. Drop-seq (analysis of snRNA-seq data, clustering, marker genes), Census-seq (estimation of chimerism in whole-genome sequencing data) and Dropulation analysis (estimation of chimerism in snRNA-seq data): https://github.com/broadinstitute/Drop-seq; alignment and variant detection of Illumina whole genome sequencing data: bwa (https://github.com/lh3/bwa), GATK (https://gatk.broadinstitute.org), BCFtools (https://github.com/samtools/bcftools), samtools (http://www.htslib.org/download), Picard Tools (https://broadinstitute.github.io/picard); R environment (https://www.rstudio.com/products/rstudio/download and https://www.r-project.org); single-cell analysis in R: Seurat (https://github.com/satijalab/seurat), cell type identification: scType (https://github.com/IanevskiAleksandr/sc-type), SCT transform (https://github.com/satijalab/sctransform/); PEER latent factor analysis (https://github.com/PMBio/peer).

## Data availability

Brain snRNA-seq of 6 marmosets (CJ022, CJ023, CJ025, CJ026, CJ027, CJ028) were generated as part of the NIH’s Brain Initiative Cell Census Network (BICCN) project, while brain snRNA-seq of 5 marmosets (CJ001, CJ006, CJ007, CJ023, CJ102), and all blood, liver and kidney snRNA-seq were generated for this project. All snRNA-seq datasets are available in the BICCN NeMO portal (https://assets.nemoarchive.org/dat-hsgdsgu and https://assets.nemoarchive.org/dat-1je0mn3). All whole genome sequencing datasets will be provided in a manuscript (in preparation) that will describe naturally occurring genome variation in captive marmosets.

## Acknowledgements

This work was supported by the NIH BRAIN Initiative U01MH114819, the Stanley Center for Psychiatric Research at the Broad Institute of MIT and Harvard, the James and Patricia Poitras Center for Psychiatric Disorders Research at MIT and the Hock E. Tan and K. Lisa Yang Center for Autism Research at MIT.

---

**Supplementary Table 1.** Number of microglia and macrophage cells identified in brain datasets and number of nuclei profiled in blood, liver and kidney.

| ID | Tissue | Microglia | Macrophage | Other Cell Types | Total nuclei | Percent Microglia | Percent Macrophage |
| --- | --- | --- | --- | --- | --- | --- | --- |
| CJ001 | Brain | 1,063 | 16 | 89,599 | 90,678 | 1.17% | 0.02% |
| CJ006 | Brain | 632 | 259 | 165,272 | 166,163 | 0.38% | 0.16% |
| CJ007 | Brain | 1,155 | 139 | 105,407 | 106,701 | 1.08% | 0.13% |
| CJ022 | Brain | 3,659 | 228 | 217,312 | 221,199 | 1.65% | 0.10% |
| CJ023 | Brain | 581 | 38 | 79,692 | 80,311 | 0.72% | 0.05% |
| CJ025 | Brain | 1,366 | 131 | 210,628 | 212,125 | 0.64% | 0.06% |
| CJ026 | Brain | 693 | 26 | 62,743 | 63,462 | 1.09% | 0.04% |
| CJ027 | Brain | 8,398 | 167 | 342,419 | 350,984 | 2.39% | 0.05% |
| CJ028 | Brain | 18,185 | 172 | 478,765 | 497,333 | 3.66% | 0.03% |
| CJ029 | Brain | 11,874 | 285 | 320,000 | 332,159 | 3.57% | 0.09% |
| CJ102 | Brain | 1,124 | 43 | 56,602 | 57,769 | 1.95% | 0.07% |
| CJ026 | Blood | NA | NA | 1,741 | 1,741 | NA | NA |
| CJ026 | Liver | NA | NA | 10,877 | 10,877 | NA | NA |
| CJ026 | Kidney | NA | NA | 9,262 | 9,262 | NA | NA |
| CJ027 | Blood | NA | NA | 2,529 | 2,529 | NA | NA |
| CJ028 | Blood | NA | NA | 10,042 | 10,042 | NA | NA |

**Supplementary Table 2.** Comparison of chimerism between CJ028’s two birth siblings, in blood and in brain myeloid cells (microglia and macrophage). *P*-values are from a two-sided test of proportions between chimerism fractions of sibling 1 and sibling 2 using the prop.test function in R.

| Cell type | Total nuclei | Sibling 1 nuclei | Sibling 2 nuclei | Sibling 1 Fraction | Sibling 2 Fraction | Sibling 1 + Sibling 2 Fraction | *P*-value |
| --- | --- | --- | --- | --- | --- | --- | --- |
| microglia and macrophage | 19701 | 6922 | 2689 | 0.35 | 0.14 | 0.49 | $<2.2 \times 10^{-16}$ |
| microglia only | 19447 | 6873 | 2621 | 0.35 | 0.13 | 0.49 | $<2.2 \times 10^{-16}$ |
| macrophage only | 254 | 49 | 68 | 0.20 | 0.27 | 0.46 | $5.8 \times 10^{-2}$ |
| blood | 10042 | 1758 | 6739 | 0.18 | 0.67 | 0.85 | $<2.2 \times 10^{-16}$ |

**Supplementary Table 3.** Summary of context versus genetic effects analysis, with two brain regions of an animal as two contexts. The analysis described in Fig. 5D-F was repeated across all animals and brain regions with at least 60 cells that are available for analysis in each context, and the summary of the correlations are tabulated here. The correlations are plotted in Supplementary Fig. 5. Abbreviations; STR: striatum; Thal: thalamus; Hippo: hippocampus; Hyp: hypothalamus; BF: basal forebrain.

| Host ID | Sibling ID | Brain Region1 | Brain Region2 | Genetic Effects Spearman Correlation | Context Effects Spearman Correlation |
| --- | --- | --- | --- | --- | --- |
| CJ022 | CJ106 | Cortex | Thal | 0.1 | 0.73 |
| CJ025 | CJ104 | Cortex | STR | 0.03 | 0.2 |
| CJ027 | CJ140 | Cortex | Thal | 0.13 | 0.64 |
| CJ027 | CJ140 | Cortex | STR | 0.14 | 0.69 |
| CJ027 | CJ140 | Cortex | Hippo | 0.08 | 0.46 |
| CJ027 | CJ140 | Cortex | Hyp | 0.03 | 0.68 |
| CJ027 | CJ140 | Thal | Hippo | 0.04 | 0.56 |
| CJ027 | CJ140 | Thal | Hyp | 0.09 | 0.66 |
| CJ027 | CJ140 | STR | Hippo | 0.08 | 0.43 |
| CJ027 | CJ140 | STR | Hyp | 0.08 | 0.59 |
| CJ027 | CJ140 | Hippo | Hyp | 0.04 | 0.51 |
| CJ028 | CJ141 | Cortex | Thal | 0.1 | 0.66 |
| CJ028 | CJ142 | Cortex | Thal | 0.03 | 0.55 |
| CJ141 | CJ142 | Cortex | Thal | 0.07 | 0.59 |
| CJ028 | CJ141 | Cortex | STR | 0.13 | 0.68 |
| CJ028 | CJ142 | Cortex | STR | 0.1 | 0.58 |
| CJ141 | CJ142 | Cortex | STR | 0.09 | 0.61 |
| CJ028 | CJ141 | Cortex | Hippo | 0.06 | 0.39 |
| CJ028 | CJ142 | Cortex | Hippo | 0 | 0.33 |
| CJ141 | CJ142 | Cortex | Hippo | 0.04 | 0.38 |
| CJ028 | CJ141 | Cortex | BF | 0.13 | 0.73 |
| CJ028 | CJ142 | Cortex | BF | 0.07 | 0.6 |
| CJ141 | CJ142 | Cortex | BF | 0.09 | 0.6 |
| CJ028 | CJ141 | Cortex | Hyp | 0.06 | 0.78 |
| CJ028 | CJ142 | Cortex | Hyp | 0.07 | 0.72 |
| CJ141 | CJ142 | Cortex | Hyp | 0.08 | 0.75 |
| CJ028 | CJ141 | Cortex | Amygdala | 0.14 | 0.65 |
| CJ028 | CJ142 | Cortex | Amygdala | 0.1 | 0.53 |
| CJ141 | CJ142 | Cortex | Amygdala | 0.12 | 0.57 |
| CJ028 | CJ141 | STR | Hippo | 0.06 | 0.63 |
| CJ028 | CJ142 | STR | Hippo | 0.01 | 0.56 |
| CJ141 | CJ142 | STR | Hippo | 0.03 | 0.61 |
| CJ028 | CJ141 | STR | BF | 0.14 | 0.63 |
| CJ028 | CJ142 | STR | BF | 0.06 | 0.5 |
| CJ141 | CJ142 | STR | BF | 0.05 | 0.52 |
| CJ028 | CJ141 | Hyp | Amygdala | 0.14 | 0.68 |
| CJ028 | CJ142 | Hyp | Amygdala | 0.21 | 0.65 |
| CJ141 | CJ142 | Hyp | Amygdala | 0.07 | 0.68 |
| CJ029 | CJ143 | Cortex | Thal | 0.09 | 0.61 |
| CJ029 | CJ143 | Cortex | STR | 0.06 | 0.5 |
| CJ029 | CJ143 | Cortex | Hippo | 0.05 | 0.62 |
| CJ029 | CJ143 | Cortex | BF | 0.09 | 0.79 |
| CJ029 | CJ143 | Cortex | Hyp | 0.07 | 0.78 |
| CJ029 | CJ143 | Cortex | Amygdala | 0.07 | 0.32 |
| CJ029 | CJ143 | Thal | STR | 0.09 | 0.48 |
| CJ029 | CJ143 | Thal | Hippo | 0.06 | 0.71 |
| CJ029 | CJ143 | Thal | BF | 0.14 | 0.75 |
| CJ029 | CJ143 | Thal | Hyp | 0.13 | 0.8 |
| CJ029 | CJ143 | Thal | Amygdala | 0.05 | 0.52 |
| CJ029 | CJ143 | Hippo | BF | 0.06 | 0.77 |
| CJ029 | CJ143 | Hippo | Hyp | 0.03 | 0.78 |
| CJ102 | CJ103 | Cortex | STR | 0.07 | 0.32 |

**Supplementary Table 4.** Clustering parameters used to identify microglia and macrophage cell types, and thresholds for identifying host-sibling doublets. The final number of microglia and macrophages after the second round of clustering are in Supplementary Table 2. A cell is assigned as a doublet if the Dropulation tool DetectDoublets assigned the highest likelihood for the cell as a doublet and if the log10 of the best likelihood minus the log10 of the second-best likelihood (lrt_test_stat, calculated by DetectDoublets) is greater than the doublet detection threshold (last column).

| Marm<br>oset<br>ID | Tissue | Resolution<br>parameter | Nearest<br>neighbor<br>parameter | UMI threshold (min<br>log-UMI per cell) | Doublet Detection<br>Threshold<br>(lrt_test_stat) |
| --- | --- | --- | --- | --- | --- |
| CJ001 | Brain | 0.8 | 10 | 1.2 | 0.0 |
| CJ006 | Brain | 0.5 | 10 | 1.2 | 5.0 |
| CJ007 | Brain | 0.5 | 20 | 1.2 | 10.0 |
| CJ022 | Brain Drop-seq | 0.5 | 20 | 1.2 | 2.0 |
| CJ022 | Brain 10X | 0.5 | 20 | 1.5 | 2.0 |
| CJ023 | Brain | 0.5 | 20 | 1.2 | 3.0 |
| CJ025 | Brain | 0.5 | 20 | 1.0 | 0.5 |
| CJ026 | Brain | 0.5 | 20 | 1.0 | 2.0 |
| CJ026 | Blood | 0.5 | 20 | 1.4 | 4.0 |
| CJ026 | Liver | 0.5 | 20 | 1.9 | 8.0 |
| CJ026 | Kidney | 0.5 | 20 | 1.9 | 8.0 |
| CJ027 | Brain Cortex A | 0.5 | 20 | 1.5 | 20.0 |
| CJ027 | Brain Cortex B | 0.5 | 20 | 1.5 | 20.0 |
| CJ027 | Brain others | 0.5 | 20 | 1.5 | 20.0 |
| CJ027 | Blood | 0.5 | 20 | 1.5 | 1.0 |
| CJ028 | Brain Cortex A | 0.5 | 20 | 1.5 | 25.0 |
| CJ028 | Brain Cortex B | 0.8 | 10 | 1.5 | 25.0 |
| CJ028 | Brain others | 0.5 | 20 | 1.5 | 25.0 |
| CJ028 | Blood | 0.5 | 20 | 1.5 | 1.0 |
| CJ029 | Brain Cortex | 0.5 | 20 | 1.5 | 20.0 |
| CJ029 | Brain others | 0.5 | 20 | 1.5 | 20.0 |
| CJ102 | Brain, Drop-seq | 0.8 | 10 | 1.2 | 3.0 |
| CJ102 | Brain, 10X | 0.5 | 20 | 1.2 | 5.0 |

**Supplementary Table 5.** Whole genome sequencing datasets used in (1) donor-of-origin assignment from snRNA-seq (Dropulation), and (2) estimating chimerism from blood whole genome sequencing (Census-seq).

| ID | Tissue/DNA source | Sequencing coverage |
| --- | --- | --- |
| CJ001 | Fibroblast culture | 36.4X |
| CJ119 (CJ001's sibling1) | Fibroblast culture | 39.7X |
| CJ120 (CJ001's sibling2) | Fibroblast culture | 36.7X |
| CJ006 (CJ007's sibling) | Fibroblast culture | 45.9X |
| CJ007 (CJ006's sibling) | Fibroblast culture | 55.8X |
| CJ022 | Fibroblast culture | 38.4X |
| CJ106 (CJ022's sibling) | Buccal swab | 36.0X |
| CJ023 | Fibroblast culture | 34.9X |
| CJ131 (CJ023's sibling1) | Fibroblast culture | 39.5X |
| CJ116 (CJ023's sibling2) | Fibroblast culture | 28.3X |
| CJ025 | Fibroblast culture | 72.7X |
| CJ104 (CJ025's sibling) | Buccal swab | 58.0X |
| CJ026 | Fibroblast culture | 74.1X |
| CJ105 (CJ026's sibling) | Buccal swab | 43.7X |
| CJ027 | Fibroblast culture | 47.5X |
| CJ027 | Blood | 46.9X |
| CJ140 (CJ027's sibling) | Fibroblast culture | 40.2X |
| CJ028 | Fibroblast culture | 44.6X |
| CJ028 | Blood | 52.6X |
| CJ141 (CJ028's sibling1) | Fibroblast culture | 43.6X |
| CJ142 (CJ028's sibling2) | Fibroblast culture | 41.2X |
| CJ029 | Fibroblast culture | 49.2X |
| CJ143 (CJ029's sibling) | Fibroblast culture | 40.8X |
| CJ102 | Brain | 32.4X |
| CJ103 (CJ102's sibling) | Buccal swab | 38.5X |

**Supplementary Figure 1.**
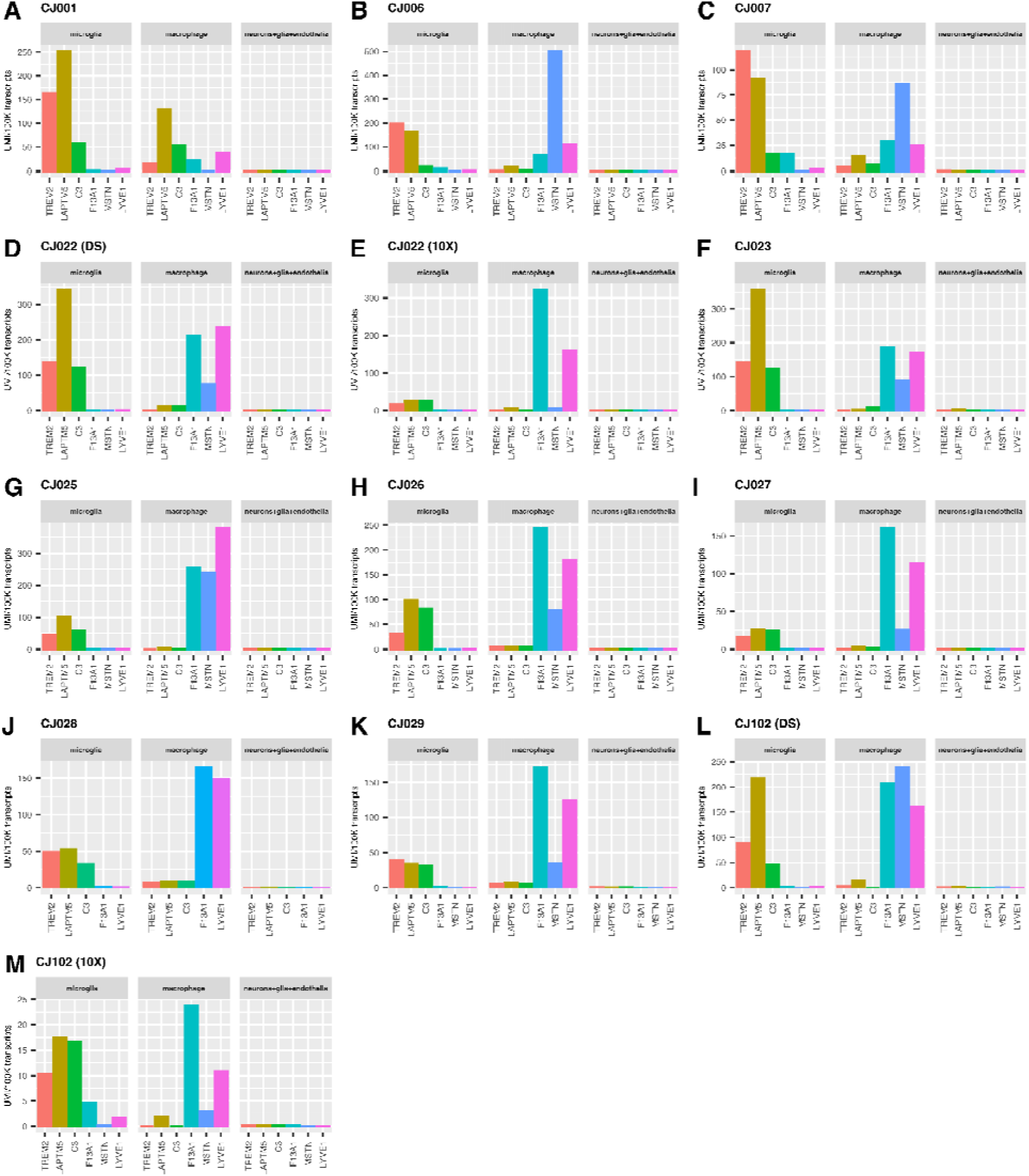
Microglia and brain macrophages can be identified in all animals. Expression of microglia and macrophage markers in microglia (left sub-panels), macrophages (middle sub-panels) and all other cell types in the brain (right sub-panels; neurons, glia (astrocytes, oligodendrocytes, polydendrocytes, ependymal cells), and endothelial cells of each animal. Marmosets CJ022 and CJ102 were profiled using two technologies (DS: Drop-seq, 10X: 10X Chromium). y-axis: unique molecular identifier (UMI, a measure of transcript abundance) of each gene across cells, summed and normalized to 100,000 transcripts. Microglia markers: *TREM2, LAPTM5, C3*; macrophage markers: *F13A1, MSTN, LYVE1*.

**Supplementary Figure 2.**
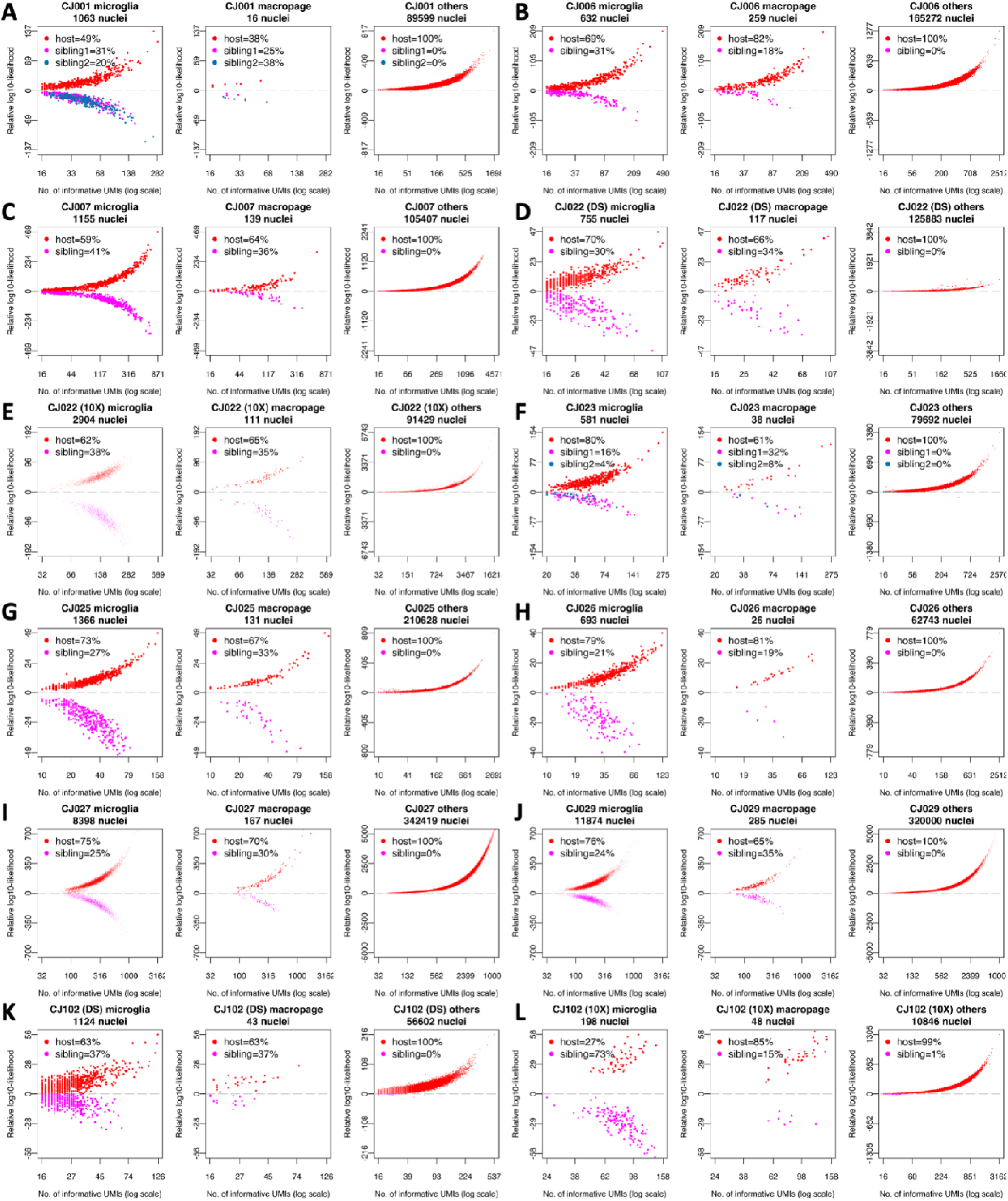
Donor-of-origin assignments from brain snRNA-seq reveals only microglia and macrophages are chimeric. (A-L) Donor of origin (Dropulation) assignments of each nucleus from brain snRNA-seq of 10 animals. Marmosets CJ022 and CJ102 were profiled using two technologies (DS: Drop-seq, 10X: 10X Chromium). For each marmoset, the snRNA-seq data are grouped into microglia, macrophage, and all other cell types (others: neurons, astrocytes, oligodendrocytes, polydendrocytes, ependymal cells, endothelial cells). x-axis: number of UMI that contains SNPs for which the host and sibling’s genomes differ, in log scale; y-axis: inferred likelihood that the cell has host genome minus likelihood that the cell has sibling genome (log10). Nuclei with positive y-values are assigned to the host and those on the negative y-axes are assigned to the sibling.

**Supplementary Figure 3.**
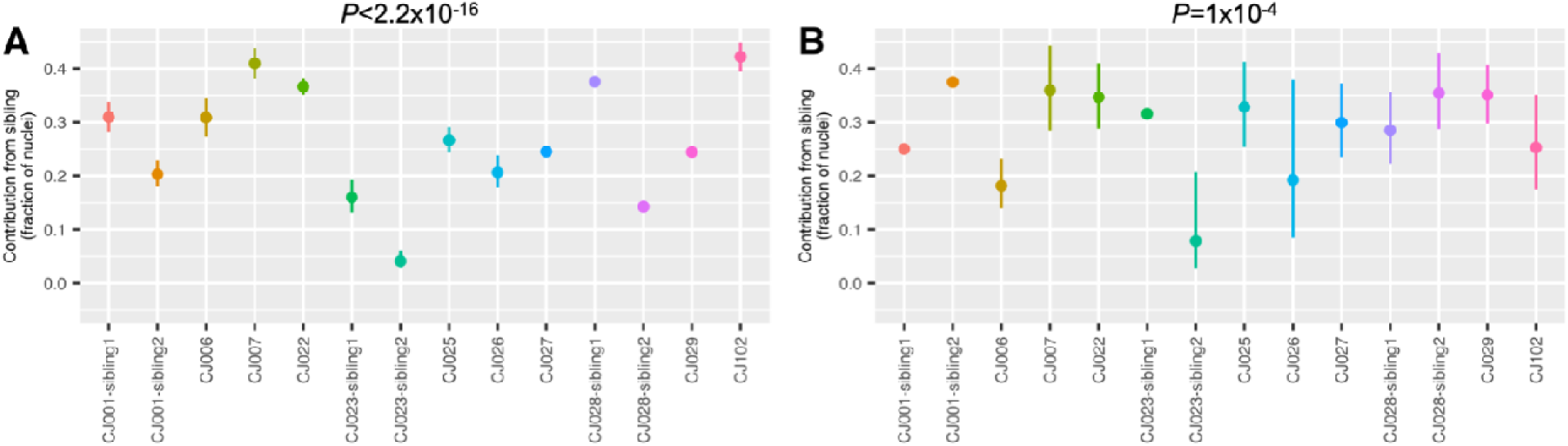
Summary of microglia (A) and macrophage (B) chimerism across animals. y-axis: fraction of twin cells. Vertical bars: binomial confidence interval (95%). *P*-values: test of heterogeneity across animals.

**Supplementary Figure 4.**
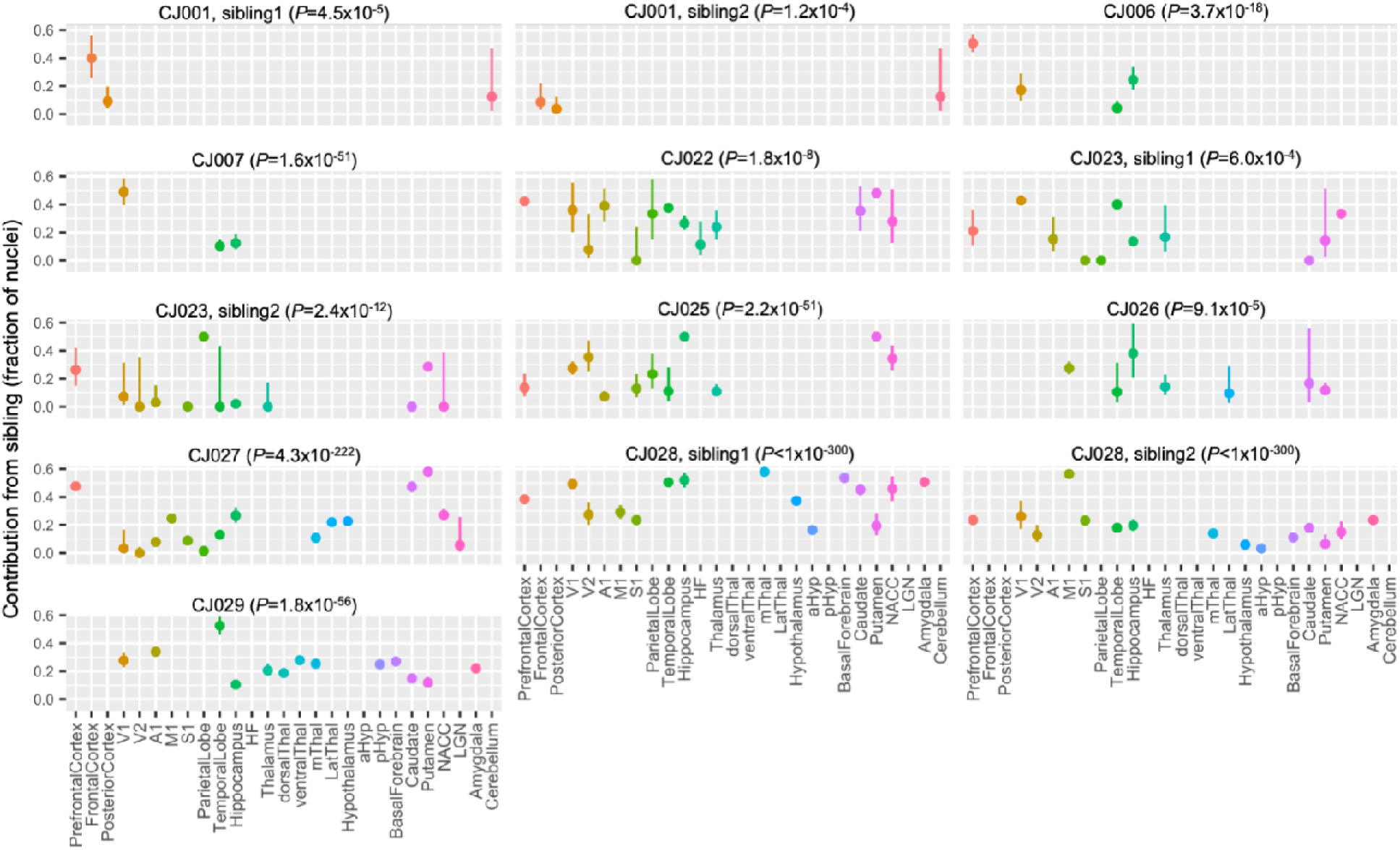
Sibling contributions to brain microglial populations vary across an animal’s brain areas. Contributions of sibling(s) to the microglial populations ascertained in finer-scale brain substructures. CJ001, CJ023 and CJ028 were part of a triplet litter and the chimerism contribution of each twin is shown in a separate panel. CJ102 was profiled in only one brain region and hence was not included in the analysis. Brain regions with missing data were not profiled in that animal. y-axis: fraction of twin cells. Vertical bars: binomial confidence interval (95%); *P*-values: test of heterogeneity across an animal’s brain regions.

**Supplementary Figure 5.**
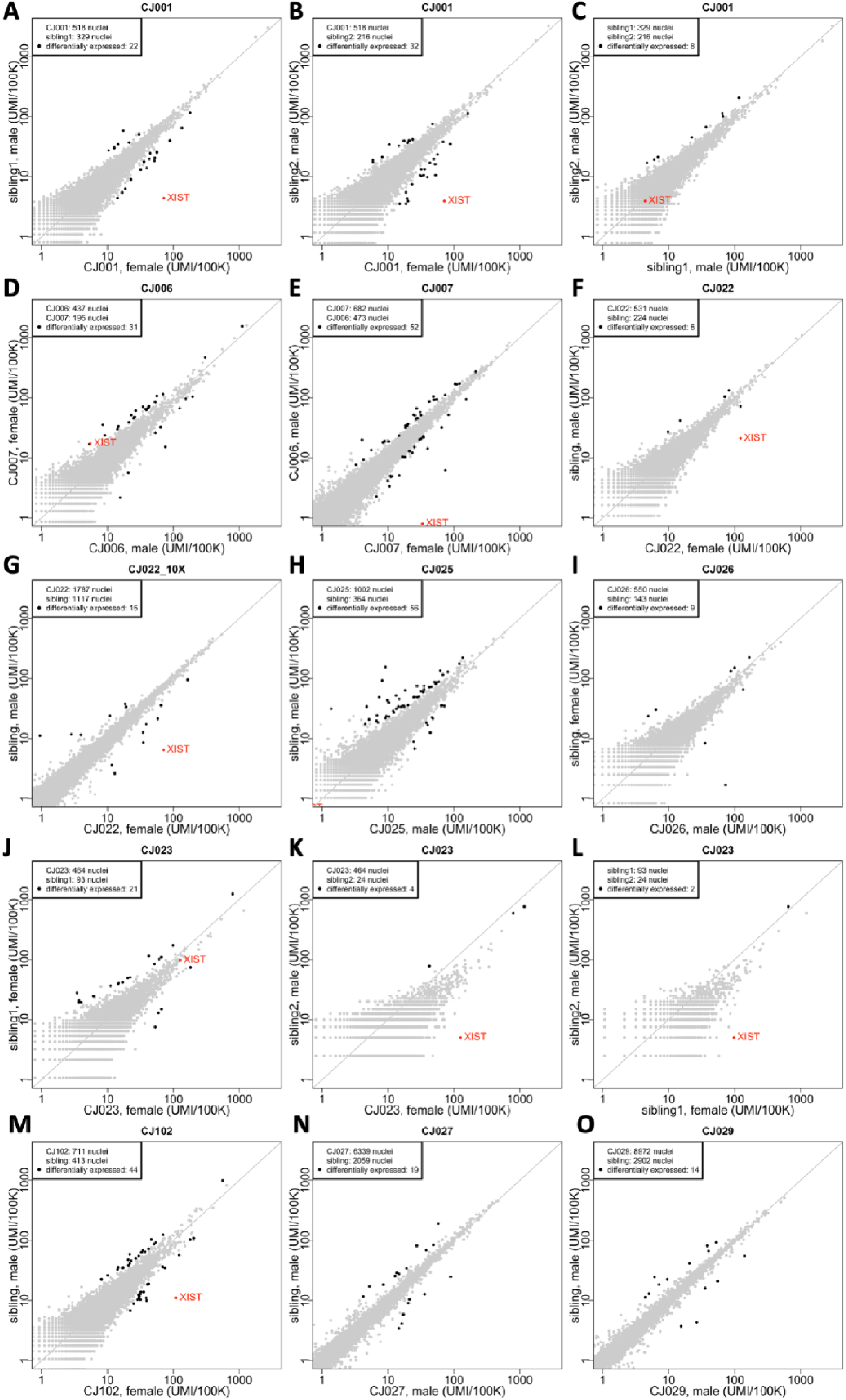
Comparison of gene expression between microglia with different genomes in each host animal’s brain. Each point represents a gene; its location on the plot represents the level of expression of that gene among microglia with two different genomes in the same animal. x– and y-axes: normalized gene expression levels (number of transcripts per 100,000 transcripts). Fold-change and *P*-values were calculated using the binomTest method from the edgeR package and differentially expressed genes (black dots) were defined as: FDR *Q*-value<0.05 and fold-change>1.5 (in either direction) and the gene must be expressed in at least 10% of at least one of the two sets of microglia being compared.

**Supplementary Figure 6.**
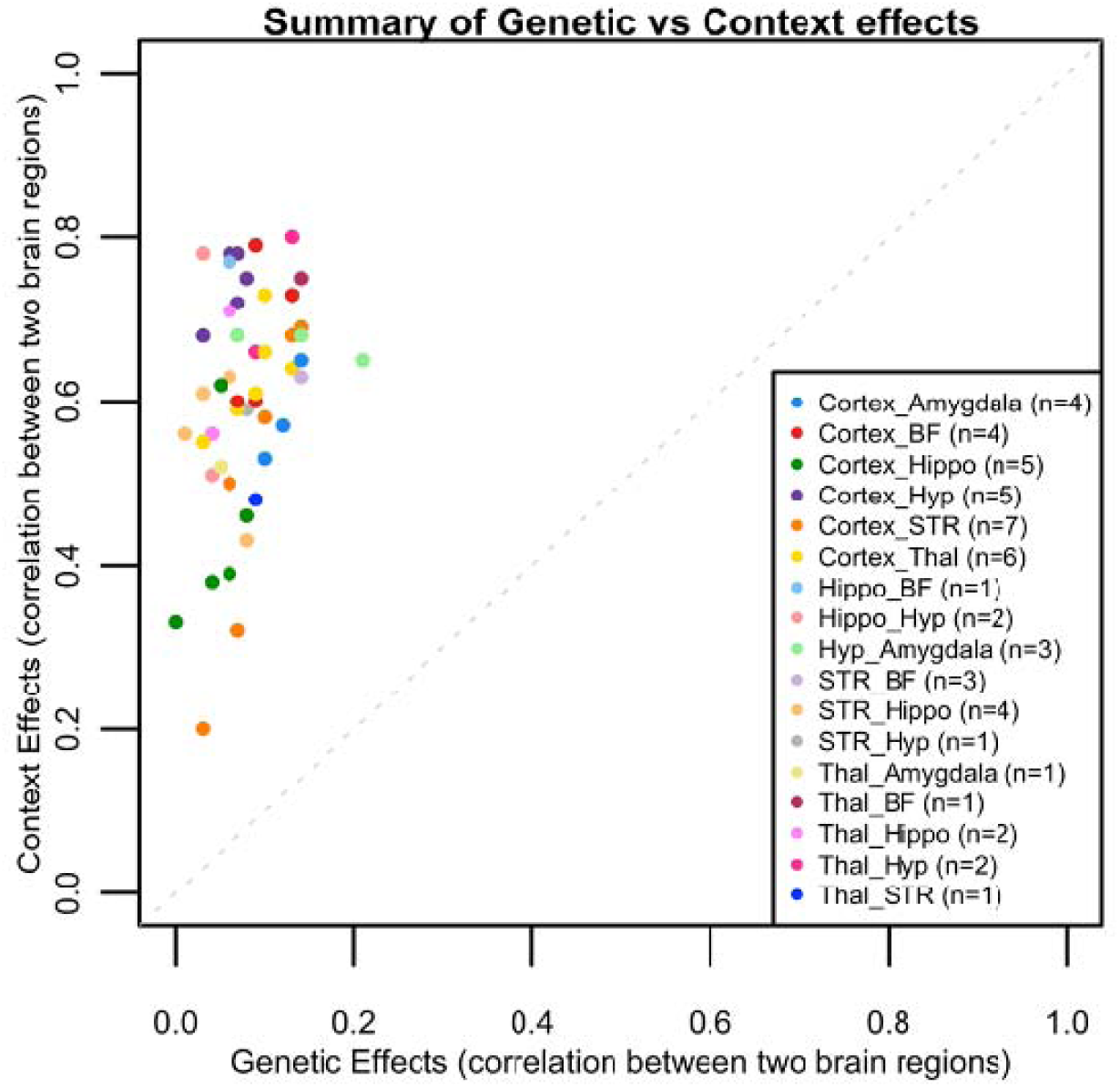
Summary of genetic versus context effects. This is a plot of the correlation values from Supplementary Table 5. Abbreviations; STR: striatum; Thal: thalamus; Hippo: hippocampus; Hyp: hypothalamus; BF: basal forebrain

**Supplementary Figure 7.**
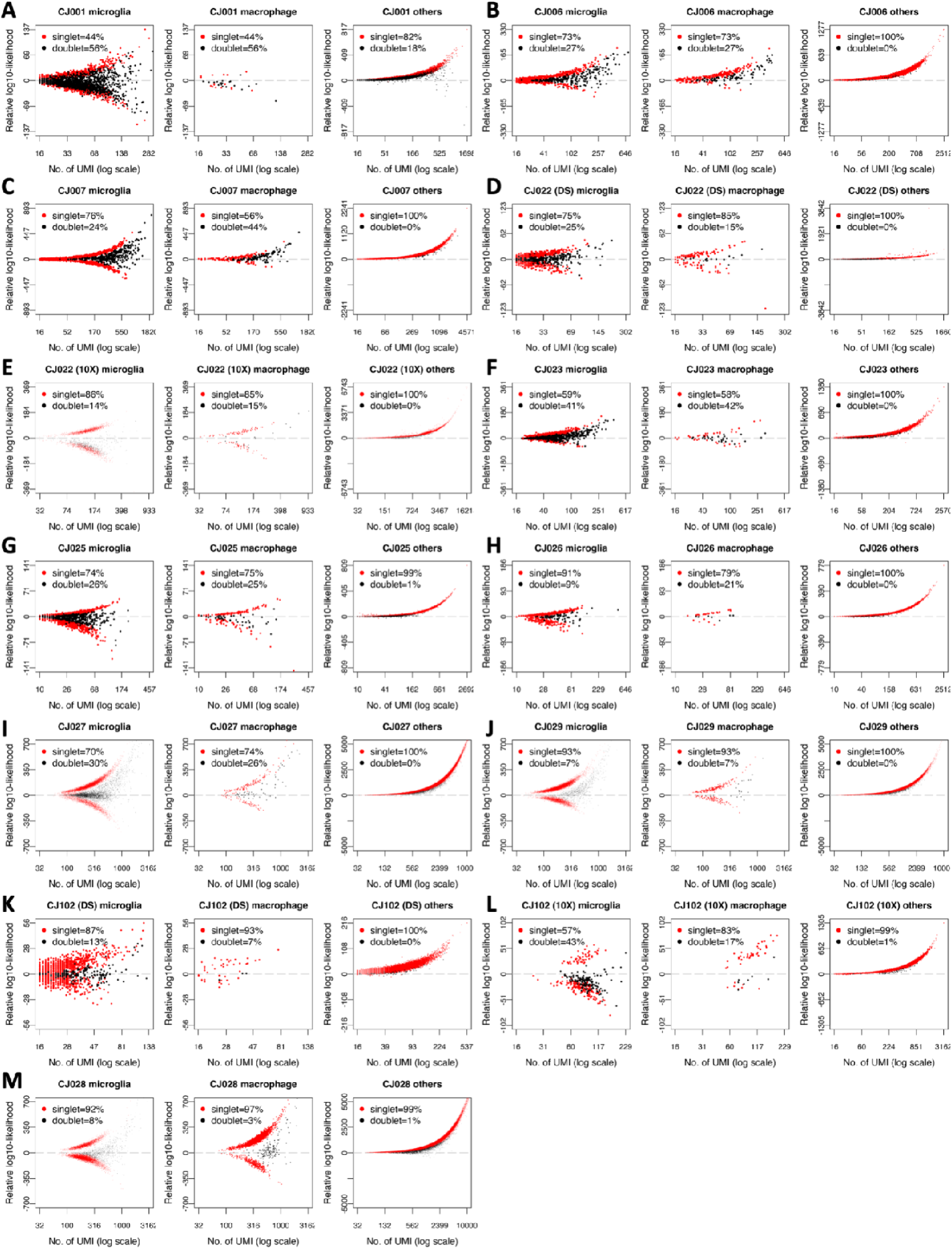
Doublet detection using host and sibling genotypes. The axes are the same as in Supplementary Fig. 2, and each dot is a nucleus. Here, nuclei that were identified as doublets and discarded in analyses were indicated (black dots). Marmosets CJ022 and CJ102 were profiled using two technologies (DS: Drop-seq, 10X: 10X Chromium).

**Supplementary Figure 8.**
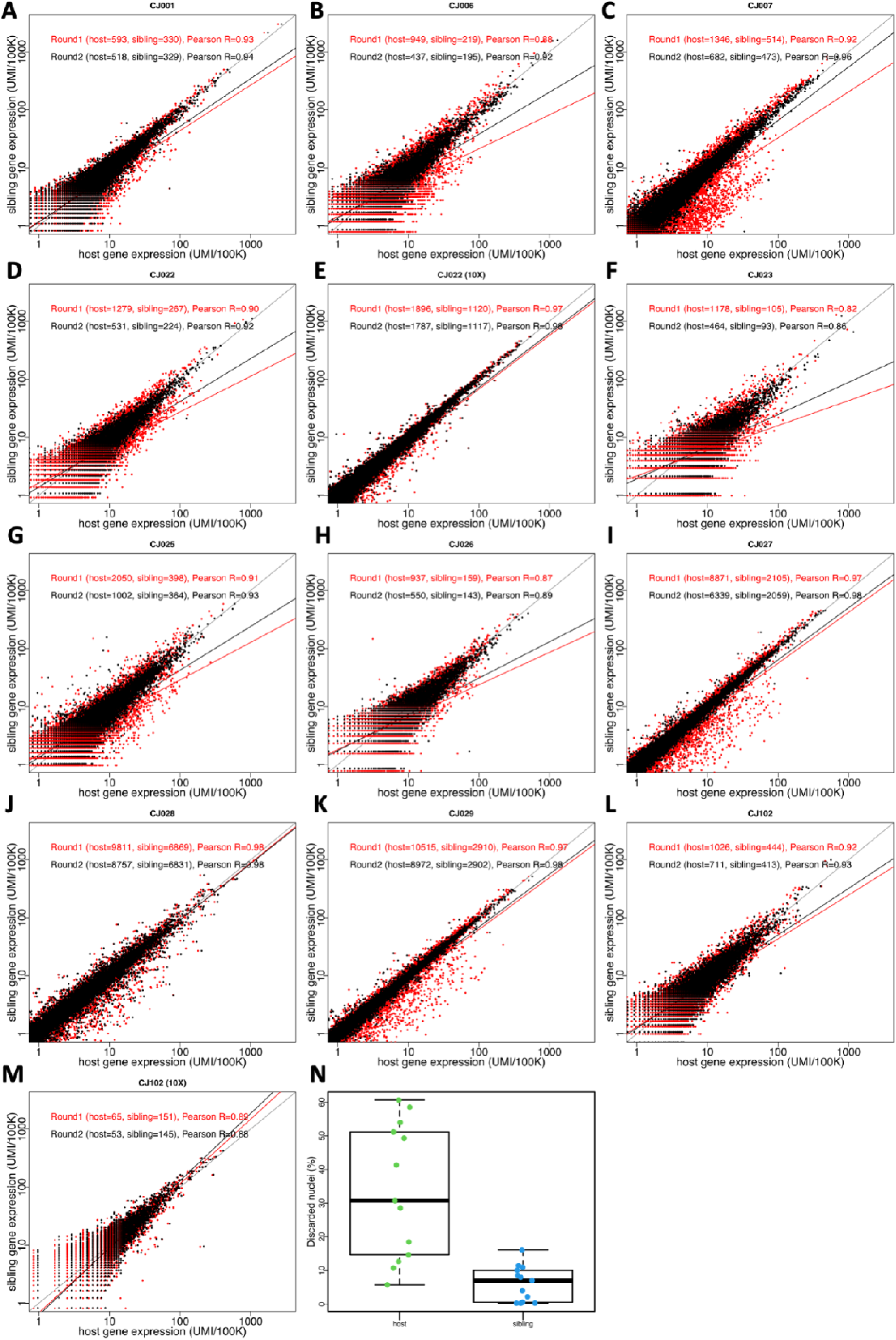
Second round clustering of microglia to discard mis-classified cells. **(A-M)** Gene expression comparison between host and sibling cells. Red dots: nuclei identified as microglia from first-round of clustering, black dots: nuclei that were retained after second-round clustering. For triplets, only the first sibling is included in the plots. Pearson correlation as calculated for each set (before and after second round clustering) and shows an improvement in correlation after discarding mis-classified cells. Marmosets CJ022 and CJ102 were profiled using two technologies (Drop-seq and 10X Chromium). (**N**) Summary (box plot) of fraction of microglia cells discarded during second round of clustering, for host and birth sibling.

**Supplementary Figure 9.**
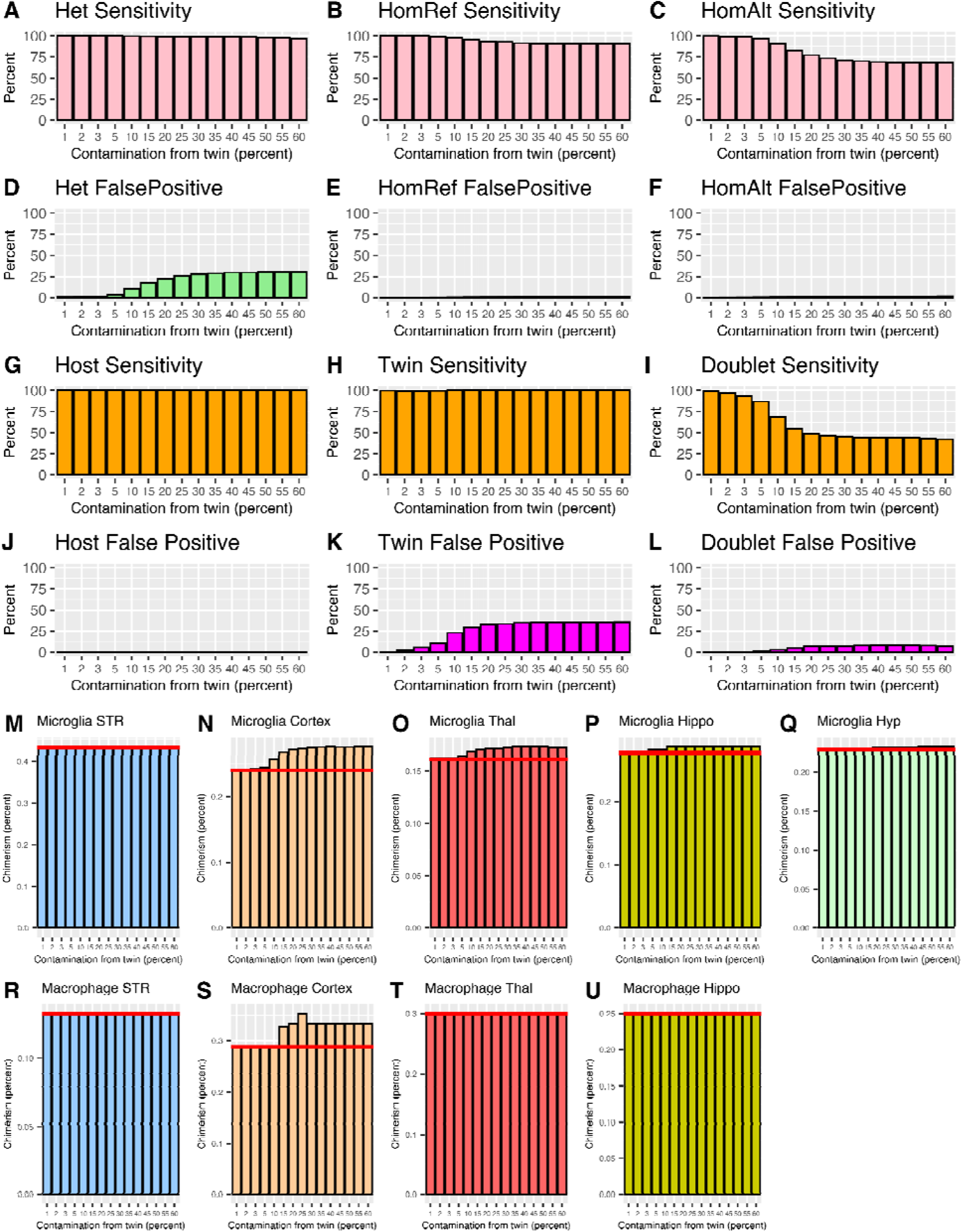
Analysis of genotyping and chimerism estimates if the whole genome sequencing data of the sibling is contaminated by the hosts’ DNA. (A)-(F) Sensitivity and false positives in genotyping; HomRef: homozygous reference, HomAlt: homozygous alternate allele, Het: heterozygous. (G)-(L) Sensitivity and false positives in donor-of-origin assignment. (M)-(Q) Microglia chimerism estimates when sibling WGS are contaminated by the hosts’ DNA, for 5 brain regions; red horizontal line: chimerism estimates when there are no errors in sibling genotypes. (R)-(U) Macrophage chimerism estimates when sibling WGS are contaminated by hosts’ DNA; red horizontal line: chimerism estimates when there are no errors in sibling genotypes.

## References

1. Aida, T., & Feng, G. (2020). The dawn of non-human primate models for neurodevelopmental disorders. Current Opinion in Genetics & Development, 65, 160–168. 10.1016/j.gde.2020.05.040

2. Choudhary, S. & Satija, R. (2022). Comparison and evaluation of statistical error models for scRNA-seq. Genome Biology, 23.1. 10.1186/s13059-021-02584-9

3. de Azevedo Fernandes NCC, Guerra JM, Díaz-Delgado J, Cunha MS, Saad LD, Iglezias SD, Ressio RA, Dos Santos Cirqueira C, Kanamura CT, Jesus IP, Maeda AY, Vasami FGS, de Carvalho J, de Araújo LJT, de Souza RP, Nogueira JS, Spinola RMF, Catão-Dias JL. (2021) Differential Yellow Fever Susceptibility in New World Nonhuman Primates, Comparison with Humans, and Implications for Surveillance. Emerg Infect Dis, Jan;27(1):47–56. doi: 10.3201/eid2701.191220

4. Feng, G., Jensen, F. E., Greely, H. T., Okano, H., Treue, S., Roberts, A. C., Fox, J. G., Caddick, S., Poo, M.-M., Newsome, W. T., & Morrison, J. H. (2020). Opportunities and limitations of genetically modified nonhuman primate models for neuroscience research. Proceedings of the National Academy of Sciences of the United States of America, 117(39), 24022–24031. 10.1073/pnas.2006515117

5. Gengozian, N., Batson, J. S., Greene, C. T., & Gosslee, D. G. (1969). Hemopoietic chimerism in imported and laboratory-bred marmosets. Transplantation, 8(5), 633–652. 10.1097/00007890-196911000-00009

6. Hammond, T. R., Robinton, D., & Stevens, B. (2018). Microglia and the Brain: Complementary Partners in Development and Disease. Annual Review of Cell and Developmental Biology, 34, 523–544. 10.1146/annurev-cellbio-100616-060509

7. Handsaker, R. E., Van Doren, V., Berman, J. R., Genovese, G., Kashin, S., Boettger, L. M., & McCarroll, S. A. (2015). Large multiallelic copy number variations in humans. Nature Genetics, 47(3), 296–303. 10.1038/ng.3200

8. Ianevski, A., Giri, A. K., & Aittokallio, T. (2022). Fully-automated and ultra-fast cell-type identification using specific marker combinations from single-cell transcriptomic data. Nature Communications, 13(1), 1246. 10.1038/s41467-022-28803-w

9. Krienen, F. M., Goldman, M., Zhang, Q., Del Rosario R, C. H., Florio, M., Machold, R., Saunders, A., Levandowski, K., Zaniewski, H., Schuman, B., Wu, C., Lutservitz, A., Mullally, C. D., Reed, N., Bien, E., Bortolin, L., Fernandez-Otero, M., Lin, J. D., Wysoker, A., … McCarroll, S. A. (2020). Innovations present in the primate interneuron repertoire. Nature, 586(7828), 262–269. 10.1038/s41586-020-2781-z

10. Krienen, F. M., Levandowski, K. M., Zaniewski, H., del Rosario, R. C. H., Schroeder, M. E., Goldman, M., Lutservitz, A., Zhang, Q., Li, K. X., Beja-Glasser, V. F., Sharma, J., Shin, T. W., Mauermann, A., Wysoker, A., Nemesh, J., Kashin, S., Vergara, J., Chelini, G., Dimidschstein, J., … Feng, G. (2023). A marmoset brain cell census reveals persistent influence of developmental origin on neurons. Sciences Advances, 9(41) DOI: 10.1126/sciadv.adk3986

11. Li, H., & Durbin, R. (2010). Fast and accurate long-read alignment with Burrows-Wheeler transform. Bioinformatics, 26(5), 589–595. 10.1093/bioinformatics/btp698

12. Ling, E., Nemesh, J., Goldman, M., Kamitaki, N., Reed, N., Handsaker, R., Genovese, G., Vogelgsang, J., Gerges, S., Kashin, S., Ghosh, S., … McCarroll S. (2024). A concerted neuron–astrocyte program declines in ageing and schizophrenia. Nature, 627, 604–611. 10.1038/s41586-024-07109-5

13. Macosko, E. Z., Basu, A., Satija, R., Nemesh, J., Shekhar, K., Goldman, M., Tirosh, I., Bialas, A. R., Kamitaki, N., Martersteck, E. M., Trombetta, J. J., Weitz, D. A., Sanes, J. R., Shalek, A. K., Regev, A., & McCarroll, S. A. (2015). Highly Parallel Genome-wide Expression Profiling of Individual Cells Using Nanoliter Droplets. Cell, 161(5), 1202–1214. 10.1016/j.cell.2015.05.002

14. McKenna, A., Hanna, M., Banks, E., Sivachenko, A., Cibulskis, K., Kernytsky, A., Garimella, K., Altshuler, D., Gabriel, S., Daly, M., & DePristo, M. A. (2010). The Genome Analysis Toolkit: a MapReduce framework for analyzing next-generation DNA sequencing data. Genome Research, 20(9), 1297–1303. 10.1101/gr.107524.110

15. Mitchell, J. M., Nemesh, J., Ghosh, S., Handsaker, R. E., Mello, C. J., Meyer, D., Raghunathan, K., de Rivera, H., Tegtmeyer, M., Hawes, D., Neumann, A., Nehme, R., Eggan, K., & McCarroll, S. A. (2020). Mapping genetic effects on cellular phenotypes with “cell villages.” bioRxiv, 2020.06.29.174383. 10.1101/2020.06.29.174383

16. Niblack, G. D., Kateley, J. R., & Gengozian, N. (1977). T-and B-lymphocyte chimerism in the marmoset. Immunology, 32(2), 257–263. https://www.ncbi.nlm.nih.gov/pubmed/139360

17. Perdiguero, E. G., & Geissmann, F. (2016). The development and maintenance of resident macrophages. Nature Immunology, 17(1), 2–8. 10.1038/ni.3341

18. Robinson, M. D., McCarthy, D. J., & Smyth, G. K. (2010). edgeR: a Bioconductor package for differential expression analysis of digital gene expression data. Bioinformatics, 26(1), 139–140. 10.1093/bioinformatics/btp616

19. Ross, C. N., French, J. A., & Orti, G. (2007). Germline chimerism and paternal care in marmosets (Callithrix kuhlii). Proceedings of the National Academy of Sciences of the United States of America, 104(15), 6278–6282. 10.1073/pnas.0607426104

20. Saunders, A., Macosko, E. Z., Wysoker, A., Goldman, M., Krienen, F. M., de Rivera, H., Bien, E., Baum, M., Bortolin, L., Wang, S., Goeva, A., Nemesh, J., Kamitaki, N., Brumbaugh, S., Kulp, D., & McCarroll, S. A. (2018). Molecular Diversity and Specializations among the Cells of the Adult Mouse Brain. Cell, 174(4), 1015–1030 e16. 10.1016/j.cell.2018.07.028

21. Schafer, D. P., Lehrman, E. K., Kautzman, A. G., Koyama, R., Mardinly, A. R., Yamasaki, R., Ransohoff, R. M., Greenberg, M. E., Barres, B. A., & Stevens, B. (2012). Microglia sculpt postnatal neural circuits in an activity and complement-dependent manner. Neuron, 74(4), 691–705. 10.1016/j.neuron.2012.03.026

22. Schafer, D. P., & Stevens, B. (2015). Microglia Function in Central Nervous System Development and Plasticity. Cold Spring Harbor Perspectives in Biology, 7(10), a020545. 10.1101/cshperspect.a020545

23. Stegle O, Parts L, Durbin R, Winn J (2010). A Bayesian Framework to Account for Complex Non-Genetic Factors in Gene Expression Levels Greatly Increases Power in eQTL Studies. PLoS Comput Biol, 6(5): e1000770. 10.1371/journal.pcbi.1000770

24. Sweeney, C. G., Curran, E., Westmoreland, S. V., Mansfield, K. G., & Vallender, E. J. (2012). Quantitative molecular assessment of chimerism across tissues in marmosets and tamarins. BMC Genomics, 13, 98. 10.1186/1471-2164-13-98

25. “The Marmoset Genome Sequencing and Analysis Consortium”, (2014). The common marmoset genome provides insight into primate biology and evolution. Nature Genetics, 46(8), 850–857. 10.1038/ng.3042

26. Wells, M. F., Nemesh, J., Ghosh, S., Mitchell, J. M., Salick, M. R., Mello, C. J., Meyer, D., Pietilainen, O., Piccioni, F., Guss, E. J., Raghunathan, K., Tegtmeyer, M., Hawes, D., Neumann, A., Worringer, K. A., Ho, D., Kommineni, S., Chan, K., Peterson, B. K., … McCarroll, S. A. (2023). Natural variation in gene expression and viral susceptibility revealed by neural progenitor cell villages. Cell Stem Cell, 30(3), 312–332.e13. 10.1016/j.stem.2023.01.010

27. Wislocki, G. B. (1939). Observations on twinning in marmosets. The American Journal of Anatomy, 64(3), 445–483. 10.1002/aja.1000640305

